# Stromal control of sympathetic rewiring in metastatic lymph nodes

**DOI:** 10.64898/2026.10.21.760843

**Authors:** Jun Li, Xiang Zhang, Guang-Hong Su, Li-Peng Hu, Qing Li, Hui Li, Rui Lu, Ting-Hong Wei, Wen-Jun Huang, Yao-Qi Zhou, Jia-Xuan Xie, Yang-Yang Wang, Tian-Shang Bao, Fei Wang, Yan-Kun Li, Shan Zhang, Chun-Jie Xu, Jian Song, Shuang-Qin Yi, Min-Juan Xu, Xiao-Wei Li, Shu-Heng Jiang, Jia Xu, Xue-Li Zhang, Dong-Xue Li, Zhi-Gang Zhang

## Abstract

Metastatic lymph nodes (LNs) are dynamically remodeled tissues, but this process has been understood mainly in immune and stromal terms. We show that tumor-driven LN remodeling also involves adaptive neural rewiring. In a mouse model of nodal metastasis, sympathetic fibers, but not sensory fibers, progressively expanded during LN remodeling and tracked with metastatic progression. Fibroblastic reticular cells (FRCs) adopted a neuro-supportive program characterized by increased hepatocyte growth factor (HGF) production, while local stromal HGF acted through MET on sympathetic neurons to stimulate neurite outgrowth. Increased substrate stiffness enhanced FRC activation and HGF production, suggesting that mechanical remodeling may reinforce the neuro-supportive stromal state. Inhibition of HGF signaling or ablation of sympathetic innervation reduced LN enlargement and nodal tumor burden. These findings identify neural plasticity as a functional component of the metastatic LN niche and reveal a stromal–mechanical–neural framework that supports metastatic LN progression.

## Introduction

Lymph nodes (LNs) are highly organized secondary lymphoid organs that integrate immune surveillance, stromal architecture, vascular function, and neural input to coordinate adaptive immunity^1–3^. In addition to their established roles in antigen presentation and lymphocyte activation, LNs are innervated by both sensory and sympathetic fibers, which can regulate local immune and vascular responses through neuropeptides and catecholamines^2,4–6^. Although neural control of LN function is increasingly appreciated, whether LN innervation itself dynamically adapts during pathological remodeling remains largely unknown^2,7,8^.

A defining feature of reactive LNs is their capacity to undergo rapid expansion^3,9,10^. This process depends on extensive stromal remodeling, particularly by fibroblastic reticular cells (FRCs), which reshape the extracellular matrix (ECM), tissue mechanics, and immune cell organization^3,9–11^. During infection, such changes are typically transient and self-limited. By contrast, tumor-draining lymph nodes (TDLNs) undergo persistent remodeling and provide a specialized microenvironment for metastatic seeding and outgrowth^1,12,13^. While immune and stromal alterations in TDLNs have been extensively studied, whether tumor-driven LN remodeling is accompanied by adaptive changes in local neural architecture, and whether such neural plasticity contributes to metastatic progression, has remained unresolved^12,13^.

Here, using a tumor-induced popliteal LN metastasis model together with time-resolved whole-organ Hydrogel-based Reinforcement of Three-Dimensional Imaging Solvent-Cleared Organs (HYBRiD) imaging^14^, single-nucleus transcriptomic profiling, and functional perturbation, we define the temporal relationship between stromal remodeling, tumor progression, and neural plasticity in TDLNs. We uncover selective remodeling of sympathetic, but not sensory, innervation and identify FRCs as a major stromal population that acquires a neuro-supportive transcriptional state during this process. Mechanistically, stromal HGF signals through MET on sympathetic neurons to promote neurite outgrowth, while matrix stiffening enhances FRC activation and HGF production. Local disruption of either HGF signaling or sympathetic innervation suppresses LN remodeling and nodal metastatic progression. These findings reveal adaptive sympathetic rewiring as a functional component of the metastatic LN niche and establish a stromal–mechanical–neural framework for tumor-induced LN remodeling.

## Results

### 1. Tumor-induced LN enlargement is accompanied by selective sympathetic nerve remodeling

To determine whether tumor-driven LN remodeling includes an adaptive neural component, we established a popliteal lymph node (popLN) metastasis model by footpad inoculation of MC38 colorectal cancer cells (**Extended Data Fig. 1a, b**). We first characterized the spatiotemporal dynamics of tumor cell infiltration and stromal remodeling using tdTomato-labeled MC38 cells, PDPN staining, and HYBRiD-based whole-organ imaging at days 0, 1, 2, 3, 5, 7, and 10 after tumor inoculation^14,15^. PDPN^+^ stromal network remodeling was evident as early as day 2, whereas tdTomato^+^ MC38 cells became readily detectable in popLNs by day 3 and progressively accumulated thereafter, reaching extensive intranodal invasion by day 10 (**Fig. 1a–c**; **Extended Data Fig. 1c–e**). In parallel, popLNs progressively enlarged during this period, with significant expansion becoming evident by day 3 (**Fig. 1d**). Together, these time-course analyses revealed an ordered remodeling process characterized by early stromal network changes, followed by overt LN enlargement and progressive intranodal tumor accumulation.

**Fig. 1.**
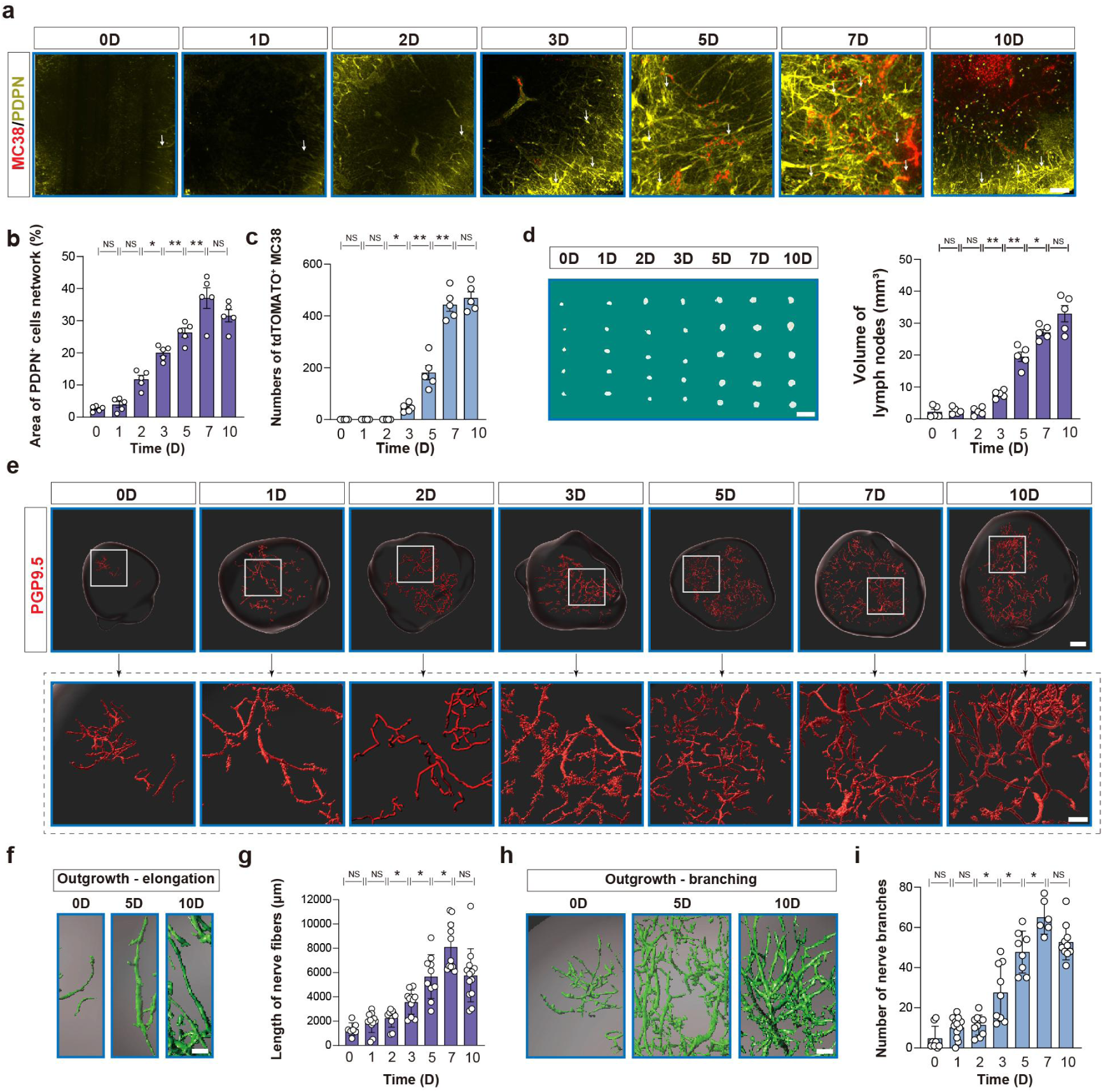
Tumor-induced LN enlargement is accompanied by adaptive intranodal nerve remodeling. a, Representative 3D images of tdTomato^+^ MC38 cells and PDPN^+^ stromal networks in popLNs at days 0, 1, 2, 3, 5, 7, and 10 after footpad tumor inoculation. Representative 3D volumes (600 μm × 600 μm × 50 μm) are shown. Scale bar, 100 μm. b, c, Quantification of the area occupied by the PDPN^+^ stromal network (b) and the number of tdTomato^+^ MC38 cells (c) in popLNs at the indicated time points. d, Representative images of popLNs collected at days 0, 1, 2, 3, 5, 7, and 10 after tumor inoculation, with quantification of popLN volume shown on the right. Scale bar, 6 mm. At least five popLNs were analyzed per group. e, Representative whole-organ 3D images of PGP9.5+ nerve fibers in popLNs at the indicated time points. Lower panels show magnified views of the boxed regions in the corresponding upper panels. Scale bars, 300 μm (upper) and 100 μm (lower). f, h, Representative reconstructions illustrating PGP9.5+ nerve fiber elongation (f) and branching (h) in popLNs at days 0, 5, and 10. Scale bars, 100 μm. g, i, Quantification of total PGP9.5^+^ nerve fiber length (g) and branch number (i) in popLNs at days 0, 1, 2, 3, 5, 7, and 10 after tumor inoculation. PGP9.5 was used as a pan-neuronal marker, and the imaging and quantification focused on intranodal parenchymal nerve fibers rather than capsular or perinodal nerve plexuses. *: *P*<0.05; **: *P*<0.01; NS: no significant difference.

We next examined intranodal innervation across the same time course. Whole-organ 3D imaging using the pan-neuronal marker PGP9.5 (UCHL1) revealed progressive increases in nerve fiber length and branching, with significant remodeling becoming evident by day 3 and increasing thereafter (**Fig. 1e–i**). Representative reconstructions further demonstrated pronounced elongation and branching of PGP9.5^+^ fibers at later stages of LN remodeling (**Fig. 1f, h**). These progressive morphological changes were consistent with active neural remodeling rather than a simple proportional increase accompanying tissue expansion.

Because LNs are innervated by both sympathetic and sensory fibers^2,4,5^, we next asked which neural subtype accounted for this remodeling. Three-dimensional imaging of TH^+^ fibers revealed a marked increase in sympathetic nerve length and branching following tumor inoculation (**Extended Data Fig. 2a-c**). To directly visualize the dynamics of sympathetic remodeling, we generated DBH-P2A-EGFP reporter mice and performed time-resolved whole-organ 3D imaging of popLNs at days 0, 1, 2, 3, 5, 7, and 10. DBH-EGFP^+^ sympathetic fibers showed progressive elongation and increased branching during tumor-induced LN remodeling, with both parameters increasing markedly as popLNs enlarged (**Fig. 2a-d**; **Extended Data Video 1; Extended Data Fig. 2d**).

**Fig. 2.**
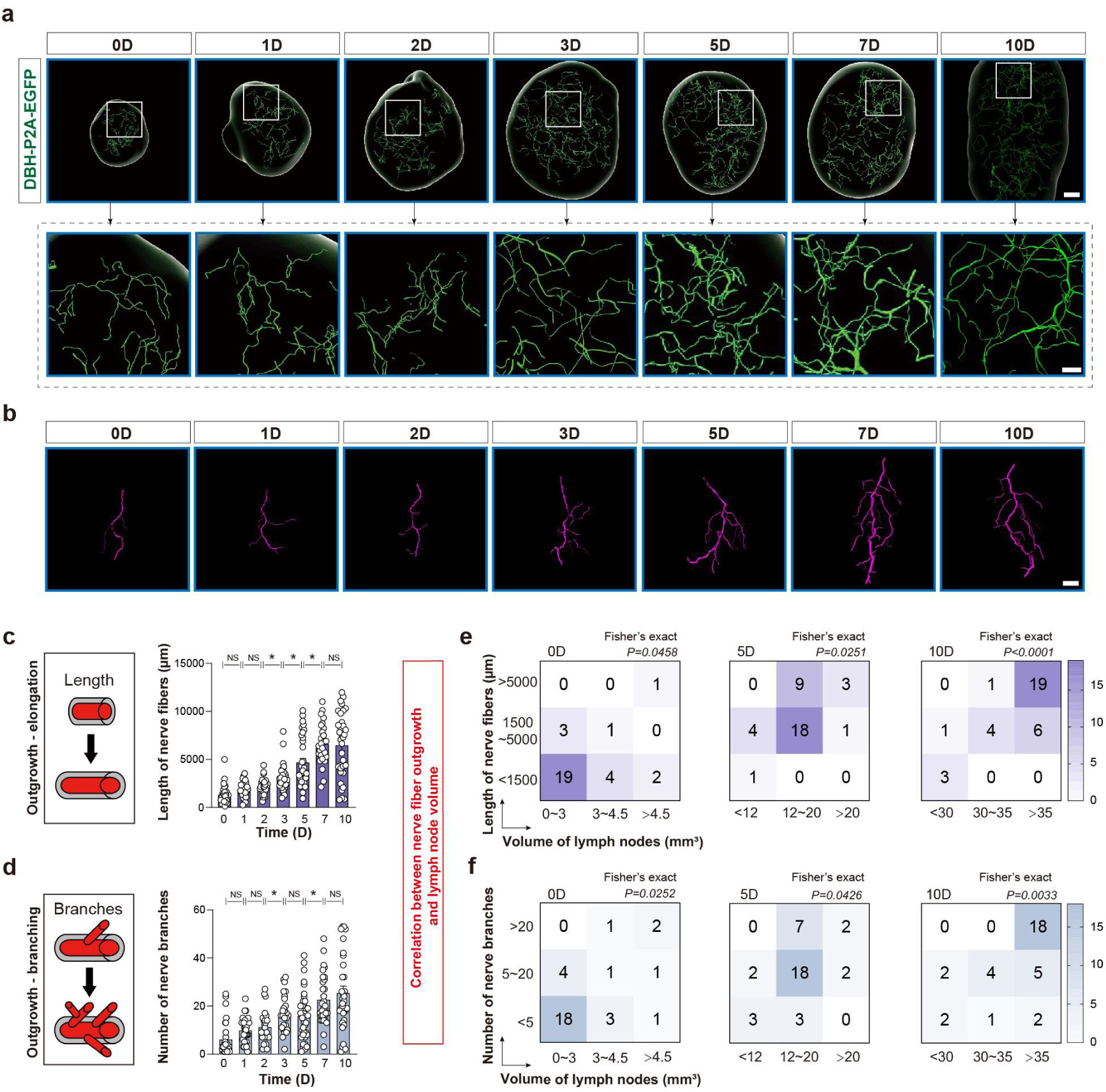
Tumor-induced LN remodeling selectively promotes sympathetic nerve outgrowth. a, Representative whole-organ 3D images of DBH-EGFP^+^ sympathetic nerve fibers in popLNs from DBH-P2A-EGFP reporter mice at days 0, 1, 2, 3, 5, 7, and 10 after tumor inoculation. Lower panels show magnified views of the boxed regions in the corresponding upper panels. Scale bars, 300 μm (upper) and 100 μm (lower). b, Representative reconstructed DBH-EGFP^+^ sympathetic nerve fibers illustrating the progressive morphological remodeling of individual fibers at the indicated time points. Scale bar, 100 μm. c, d, Quantification of DBH-EGFP^+^ sympathetic nerve fiber length (c) and branch number (d) in popLNs at the indicated time points. e, f, Association between popLN volume and sympathetic nerve remodeling. Samples were stratified according to popLN volume and DBH-EGFP^+^ sympathetic nerve fiber length (e) or branch number (f). Numbers within each cell indicate the number of samples in the corresponding category. Statistical associations were assessed using Fisher’s exact test. *: *P*<0.05; **: *P*<0.01; NS: no significant difference.

We next examined the relationship between sympathetic remodeling and LN expansion. Joint analysis of sympathetic fiber length or branch number with popLN volume revealed a progressive shift toward states characterized by both increased innervation and increased LN size, supporting a close association between sympathetic structural remodeling and LN expansion (**Fig. 2e, f**). In contrast, CALCA-P2A-BFP reporter mice showed no significant sensory nerve outgrowth at the examined time points (**Extended Data Fig. 3a–e**). Co-staining for NFL and TH confirmed the sympathetic identity of the TH^+^ nerve fibers (**Extended Data Fig. 4a, b**).

Together, these findings demonstrate that tumor-induced LN remodeling is accompanied by selective and progressive sympathetic nerve outgrowth, rather than a generalized expansion of peripheral innervation.

### 2. FRCs acquire a neuro-supportive transcriptional state during tumor-induced LN remodeling

To define the cellular programs associated with sympathetic remodeling, we performed temporal single-nucleus RNA sequencing (snRNA-seq) of popLNs at days 0, 1, 2, 3, 5, 7, and 10 after tumor inoculation, spanning early stromal remodeling through progressive nodal tumor invasion (**Fig. 3a**).

**Fig. 3.**
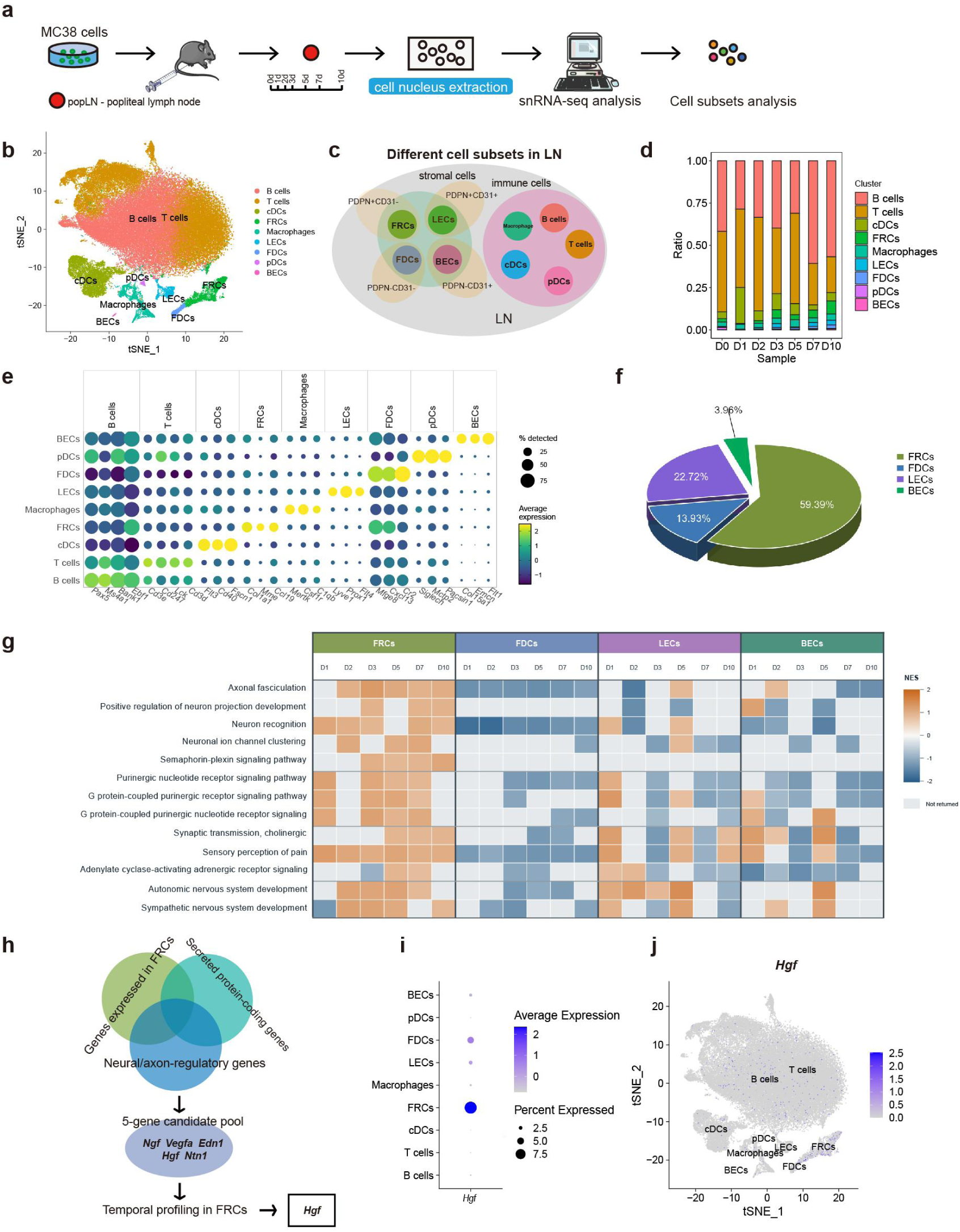
FRCs acquire a neuro-supportive state during tumor-induced LN remodeling. a, Schematic of the temporal snRNA-seq workflow. popLNs were collected at days 0, 1, 2, 3, 5, 7, and 10 after footpad inoculation of MC38 cells for nuclei isolation, sequencing, and cell population analysis. b, t-SNE visualization of the major cell populations identified by snRNA-seq. The dataset has been deposited in the Gene Expression Omnibus under accession number GSE280461. c, Schematic representation of the nine major cell populations identified in popLNs. d, Relative proportions of FRCs, FDCs, LECs, BECs, B cells, T cells, macrophages, cDCs, and pDCs at the indicated time points after tumor inoculation. e, Dot plot showing representative marker genes used to annotate the major cell populations. Dot size indicates the percentage of cells expressing each gene, and color intensity indicates average expression. f, Relative proportions of FRCs, FDCs, LECs, and BECs within the stromal compartment. g, Heatmap showing temporal enrichment of selected GO biological process terms in FRCs, FDCs, LECs, and BECs during tumor-induced LN remodeling. Enrichment at days 1, 2, 3, 5, 7, and 10 was calculated relative to day 0. h, Strategy for identifying candidate FRC-derived neuro-supportive factors by integrating neuron/axon-related genes, secreted protein-coding factors, and temporal gene expression profiles in FRCs. i, Dot plot showing *Hgf* expression across the major popLN cell populations. Dot size indicates the percentage of *Hgf*-expressing cells, and color intensity indicates average expression. j, t-SNE visualization of *Hgf* expression across popLN cell populations.

snRNA-seq resolved nine major cell populations, including FRCs, follicular dendritic cells (FDCs), lymphatic endothelial cells (LECs), blood endothelial cells (BECs), plasmacytoid dendritic cells (pDCs), conventional dendritic cells (cDCs), and multiple immune subsets (**Fig. 3b-e**; **Extended Data Fig. 5a, b; Extended Data Fig. 6a**). Cell populations were annotated according to established lineage markers and current lymphoid stromal cell nomenclature^16,17^. Time-course analysis revealed an increased relative representation of FRCs within the stromal compartment during tumor-induced LN remodeling (**Extended Data Fig. 6b, c**). In comparison, nerve-related transcriptional programs were less prominently enriched in the major immune cell populations analyzed (**Extended Data Fig. 5c**), although flow cytometry revealed substantial changes in B cells, neutrophils, and macrophages during LN remodeling (**Extended Data Fig. 7a-i**).

Within the stromal compartment, FRCs represented the largest population, accounting for approximately 59% of stromal cells (**Fig. 3f**). Time-course GO-BP analysis revealed dynamic enrichment of neuro-supportive programs in FRCs during LN remodeling, including axonal fasciculation, positive regulation of neuron projection development, and neuron recognition (**Fig. 3g**). Consistent with these findings, complementary GO and GSEA analyses showed enrichment of pathways related to autonomic and sympathetic nervous system development in FRCs (**Extended Data Fig. 8a-d**). Together, these transcriptomic analyses indicate that FRCs acquire a neuro-supportive state during tumor-induced LN remodeling.

We next sought to identify candidate FRC-derived signals that could mediate communication with remodeling nerves. By integrating genes associated with neuronal projection and axonal regulation, secreted protein-coding factors, and their temporal expression profiles in FRCs, *Hgf* emerged as a prominent candidate (**Fig. 3h**). Among the major popLN cell populations analyzed, *Hgf* expression was enriched predominantly within the FRC compartment (**Fig. 3i, j**; **Extended Data Fig. 8e; Extended Data Fig. 9a-g**). These findings nominated FRC-derived HGF as a candidate molecular link between stromal activation and sympathetic nerve remodeling.

### 3. Tumor- and matrix-activated FRCs promote neurite outgrowth through HGF

Having identified HGF as a candidate FRC-derived neuro-supportive factor, we next asked how tumor-associated stromal activation regulates HGF production and whether this signal functionally promotes neurite outgrowth. Exposure of primary LN-derived FRCs to MC38-conditioned medium induced an activated phenotype, characterized by increased *Acta2* and *Col1a1* expression and enhanced α-SMA staining (**Fig. 4a**; **Extended Data Fig. 10a–d**). Importantly, MC38-conditioned medium also increased *Hgf* expression and HGF secretion by FRCs (**Fig. 4b**), indicating that tumor-derived soluble signals are sufficient to induce an HGF-producing stromal state.

**Fig. 4.**
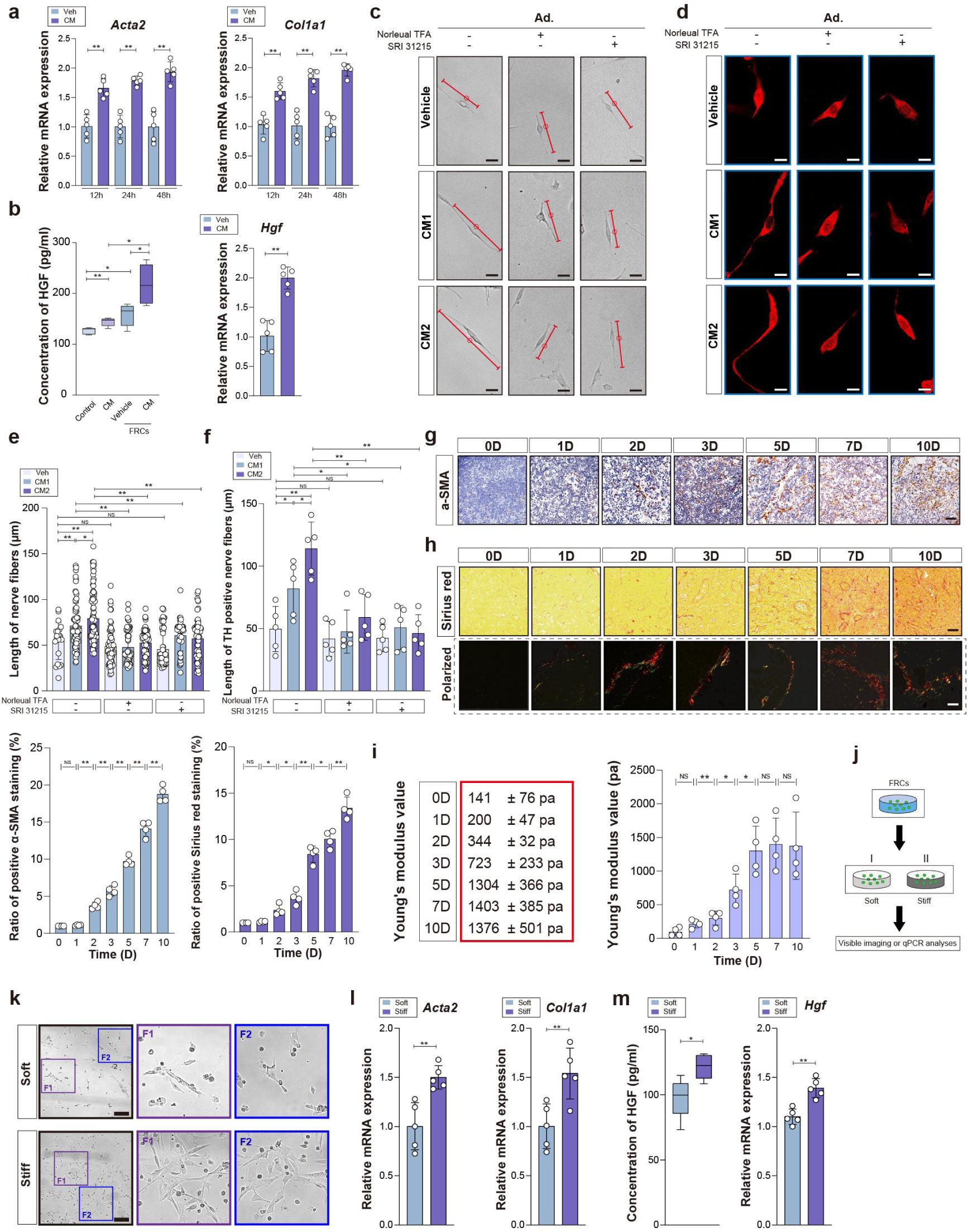
Tumor- and matrix-activated FRCs promote neurite outgrowth through HGF. a, Relative mRNA expression of Acta2 and Col1a1 in primary LN-derived FRCs treated with MC38-conditioned medium or control medium for 12, 24, and 48 h. b, HGF secretion measured by ELISA and *Hgf* mRNA expression measured by qPCR in primary FRCs treated with MC38-conditioned medium or control medium for 48 h. c, d, Representative images of PC12 cells exposed to the indicated FRC-conditioned media in the presence or absence of Norleual TFA or SRI 31215 (CM1: conditioned medium from FRCs alone; CM2: conditioned medium from FRCs exposed to MC38-conditioned medium). Scale bars, 20 μm (c) and 12 μm (d). e, f, Quantification of total neurite length (e) and TH^+^ neurite length (f) under the indicated conditions. g, Representative α-SMA immunostaining of the popLN medullary region at days 0, 1, 2, 3, 5, 7, and 10 after tumor inoculation, with corresponding quantification. h, Sirius red staining of the popLN medullary region at the indicated time points. Corresponding polarized-light images are shown below to visualize collagen fiber organization, with quantification of Sirius red staining. i, Young’s modulus of popLNs measured by atomic force microscopy at the indicated time points after tumor inoculation. j, Schematic of primary FRC culture on matrices of defined stiffness (0.5 kPa, soft; 12 kPa, stiff). k, Representative images of primary FRCs cultured on soft or stiff matrices. Scale bar, 30 μm. l, Relative mRNA expression of Acta2 and Col1a1 in primary FRCs cultured on soft or stiff matrices. m, HGF secretion measured by ELISA and *Hgf* mRNA expression measured by qPCR in primary FRCs cultured on soft or stiff matrices. *: *P*<0.05; **: *P*<0.01; NS: no significant difference.

We next tested whether this activated FRC state could promote neurite outgrowth using PC12 cells, a catecholaminergic sympathetic neuron-like model. PC12 cells were treated with conditioned medium from FRCs cultured alone (CM1) or from FRCs previously exposed to MC38-conditioned medium (CM2). Both CM1 and CM2 promoted neurite extension relative to control medium, with CM2 producing a stronger response, consistent with enhanced neuro-supportive activity following tumor-induced FRC activation (**Extended Data Fig. 10e–h**). Pharmacological inhibition of HGF signaling using Norleual TFA or SRI 31215 significantly attenuated CM-induced increases in both total neurite length and TH^+^ neurite length (**Fig. 4c–f**). These findings support HGF as a functional mediator of FRC-driven neurite outgrowth.

We then investigated whether biomechanical remodeling of the LN microenvironment further reinforces this neuro-supportive FRC program. α-SMA staining revealed stromal activation beginning during the early phase of remodeling, while Sirius red staining demonstrated progressive collagen deposition and organization during tumor progression (**Fig. 4g, h**). Atomic force microscopy showed a progressive increase in Young’s modulus from day 0 to day 10, demonstrating that tumor-induced LN remodeling is accompanied by increasing matrix stiffness (**Fig. 4i**).

To directly test whether matrix stiffness regulates the FRC neuro-supportive program, primary FRCs were cultured on matrices of defined stiffness (0.5 kPa versus 12 kPa; **Fig. 4j, k**). FRCs cultured on the stiffer matrix exhibited increased *Acta2* and *Col1a1* expression together with increased *Hgf* expression and HGF secretion (**Fig. 4l, m**). Thus, increased matrix stiffness potentiates FRC activation and HGF production, providing a biomechanical mechanism that can amplify the neuro-supportive stromal program during tumor-induced LN remodeling.

### 4. Local stromal HGF promotes sympathetic remodeling and metastatic LN progression

To determine whether HGF signaling is functionally involved in sympathetic remodeling *in vivo*, we locally inhibited HGF using either Norleual TFA or AAV-mediated *Hgf* silencing in the popliteal fat pad before MC38 inoculation (**Fig. 5a**). popLNs were analyzed at day 7, when LN enlargement and sympathetic nerve remodeling were prominent. Whole-organ 3D imaging showed that both pharmacological HGF inhibition and AAV-mediated *Hgf* silencing markedly reduced the length and branching of TH^+^ sympathetic fibers within popLNs (**Fig. 5b, c**). Similar effects were observed in DBH-P2A-EGFP reporter mice (**Fig. 5d, e**), independently confirming the suppression of sympathetic nerve remodeling following local HGF inhibition.

**Fig. 5.**
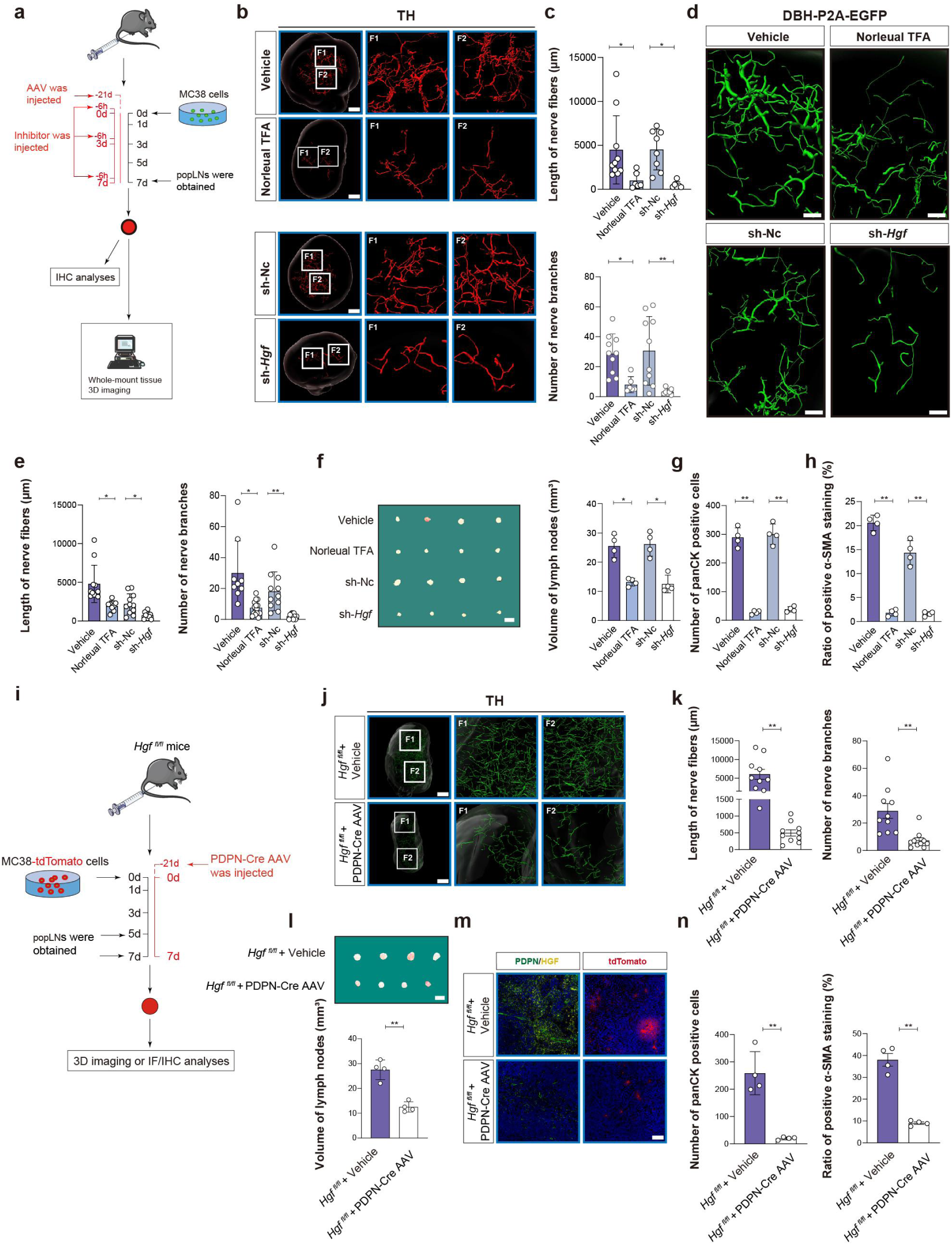
Local stromal HGF promotes sympathetic remodeling and metastatic LN progression. a, Schematic of local pharmacological or AAV-mediated HGF inhibition. C57BL/6 mice received local administration of Norleual TFA, AAV-sh-*Hgf*, or the corresponding controls in the popliteal fat pad, followed by footpad inoculation of MC38 cells. popLNs were analyzed at the indicated endpoint. b, c, Representative whole-organ 3D images of TH^+^ sympathetic nerve fibers in popLNs from mice treated with Norleual TFA, AAV-sh-*Hgf*, or the corresponding controls (b), with quantification of sympathetic nerve fiber length and branch number (c). Scale bar, 300 μm. d, e, Representative 3D images of DBH-EGFP^+^ sympathetic nerve fibers in popLNs from DBH-P2A-EGFP reporter mice under the indicated treatments (d), with quantification of nerve fiber length and branch number (e). Scale bar, 100 μm. f, Representative images and quantification of popLN volume following the indicated treatments. Scale bar, 6 mm. g, h, Quantification of panCK (g) and α-SMA (h) immunohistochemical staining in popLNs following local HGF inhibition. i, Schematic of local PDPN-Cre AAV-mediated *Hgf* deletion in *Hgf*^fl/fl^ mice. PDPN-Cre AAV or vehicle was locally delivered to the popLN region before footpad inoculation of MC38-tdTomato cells. j, k, Representative whole-organ 3D images of TH^+^ sympathetic nerve fibers in popLNs from *Hgf*^fl/fl^ mice treated with PDPN-Cre AAV or vehicle (j), with quantification of nerve fiber length and branch number (k). Scale bar, 300 μm. l, Representative images and quantification of popLN volume in *Hgf*^fl/fl^ mice following local PDPN-Cre AAV-mediated *Hgf* deletion. Scale bar, 6 mm. m, Representative immunofluorescence images showing PDPN and HGF expression, together with tdTomato^+^ MC38 cells, in popLNs from *Hgf*^fl/fl^ mice treated with PDPN-Cre AAV or vehicle. Scale bar, 100 μm. n, Quantification of panCK and α-SMA immunohistochemical staining in popLNs from *Hgf*^fl/fl^ mice following local PDPN-Cre AAV-mediated *Hgf* deletion. *: *P*<0.05; **: *P*<0.01; NS: no significant difference.

We then examined whether local HGF inhibition also affected tumor-induced LN remodeling and nodal metastatic progression. Both Norleual TFA treatment and AAV-mediated *Hgf* silencing reduced popLN enlargement following tumor challenge (**Fig. 5f**). In parallel, panCK and α-SMA analyses showed reduced nodal tumor burden and stromal activation after HGF inhibition (**Fig. 5g, h**; **Extended Data Fig. 11a, b**). Importantly, neither local Norleual TFA treatment nor AAV-mediated Hgf silencing significantly altered primary footpad tumor burden at the examined time points (**Extended Data Fig. 14a, b**), indicating that the reduced nodal metastatic burden was unlikely to be simply secondary to suppression of primary tumor growth. Together, these findings link local HGF signaling to sympathetic nerve remodeling, LN expansion, and nodal metastatic progression.

To directly assess the contribution of stromal HGF within the popLN, we used *Hgf*^fl/fl^ mice and locally delivered PDPN-Cre AAV to the popLN region to target PDPN^+^ stromal cells (**Fig. 5i**; **Extended Data Fig. 11e)**. Immunofluorescence confirmed a marked reduction of HGF within PDPN^+^ stromal regions following PDPN-Cre AAV delivery (**Fig. 5m**). Local PDPN^+^ stromal-cell-targeted deletion of *Hgf* recapitulated the effects of HGF inhibition, markedly suppressing TH^+^ sympathetic fiber elongation and branching (**Fig. 5j, k**), reducing popLN enlargement (**Fig. 5l**), and decreasing nodal metastatic burden and stromal activation, as assessed by tdTomato, panCK, and α-SMA analyses (**Fig. 5m, n**; **Extended Data Fig. 11c, d**).

Together, these findings support a functional role for local stromal HGF in sympathetic nerve remodeling and metastatic LN progression.

### 5. HGF directly promotes sympathetic neurite outgrowth through MET

To define how HGF acts on sympathetic nerves, we first examined expression of its receptor MET in popLN tissues. Immunofluorescence revealed MET expression along PGP9.5^+^ and TH^+^ nerve fibers within popLNs (**Fig. 6a**). We next isolated primary sympathetic neurons from DBH-P2A-EGFP reporter mice or C57BL/6J mice and confirmed their sympathetic identity by DBH-EGFP and TH expression. MET was readily detected in these primary sympathetic neurons by immunofluorescence (**Fig. 6b–d**), establishing that sympathetic neurons express the HGF receptor.

**Fig. 6.**
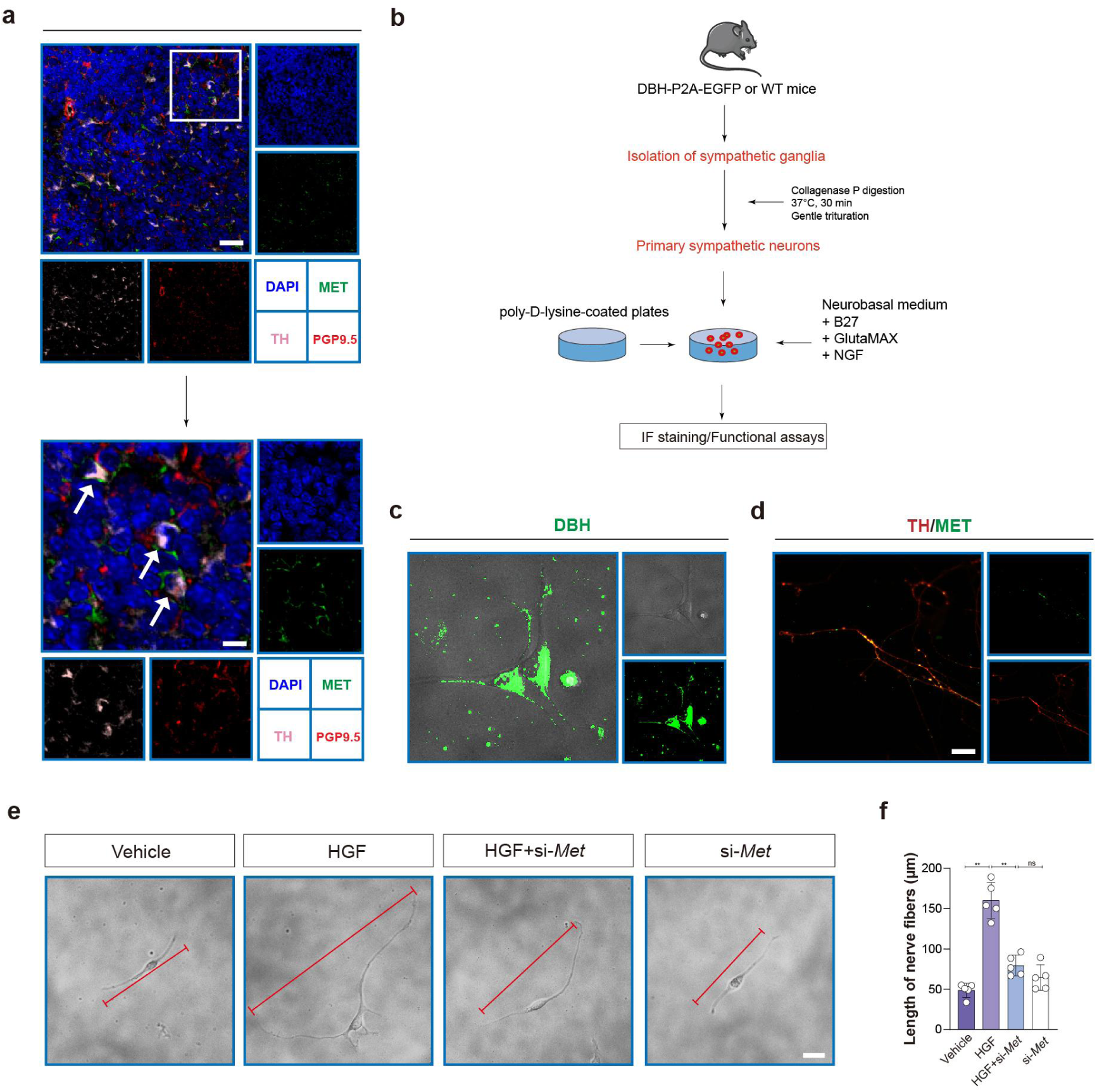
HGF promotes sympathetic neurite outgrowth through MET. a, Representative immunofluorescence images showing MET expression on PGP9.5+ and TH+ nerve fibers in popLN tissue. Boxed regions are shown at higher magnification. Scale bars, 100 μm (upper) and 30 μm (lower). b, Schematic of the isolation, culture, and functional analysis of primary sympathetic neurons from DBH-P2A-EGFP reporter mice. c, Representative images of primary DBH-EGFP+ sympathetic neurons. Scale bar, 10 μm. d, Representative immunofluorescence images showing TH and MET expression in primary sympathetic neurons. Scale bar, 10 μm. e, Representative images of primary sympathetic neurons treated with vehicle, recombinant HGF, recombinant HGF together with siRNA-mediated Met knockdown, or Met siRNA alone. Red lines indicate neurites used for length quantification. Scale bar, 20 μm. f, Quantification of neurite length in primary sympathetic neurons under the indicated conditions. *: *P*<0.05; **: *P*<0.01; NS: no significant difference.

We next tested whether HGF-induced sympathetic neurite outgrowth is MET-dependent. Recombinant HGF markedly increased neurite length in primary sympathetic neurons, whereas siRNA-mediated Met knockdown significantly attenuated this response (**Fig. 6e, f**). Together, these findings demonstrate that HGF can act directly on sympathetic neurons through MET to promote neurite outgrowth, providing a cellular mechanism linking stromal HGF to sympathetic nerve remodeling in tumor-draining LNs.

### 6. Sympathetic innervation functionally promotes metastatic LN remodeling

Having established a functional stromal HGF–MET pathway promoting sympathetic neurite outgrowth, we next asked whether sympathetic remodeling itself functionally contributes to tumor-induced LN remodeling and nodal metastasis. We first performed local chemical sympathectomy by injecting 6-hydroxydopamine (6-OHDA) into the fat pad surrounding the popLN five days before tumor inoculation (**Fig. 7a**). Three-dimensional imaging confirmed marked depletion of local sympathetic innervation following 6-OHDA treatment (**Fig. 7d**). Local sympathetic denervation significantly reduced popLN enlargement at both day 5 and day 7 after MC38 challenge (**Fig. 7b, c**). Imaging of tdTomato^+^ tumor cells and panCK staining further demonstrated a marked reduction in nodal metastatic burden following local 6-OHDA-mediated denervation (**Fig. 7e, f**). Immunohistochemical analysis additionally revealed reductions in B220^+^ B cells, CD11b^+^ myeloid cells, and Ly6G^+^ neutrophils after denervation (**Extended Data Fig. 12a, b**), indicating that sympathetic input contributes to local immune remodeling within tumor-draining LNs.

**Fig. 7.**
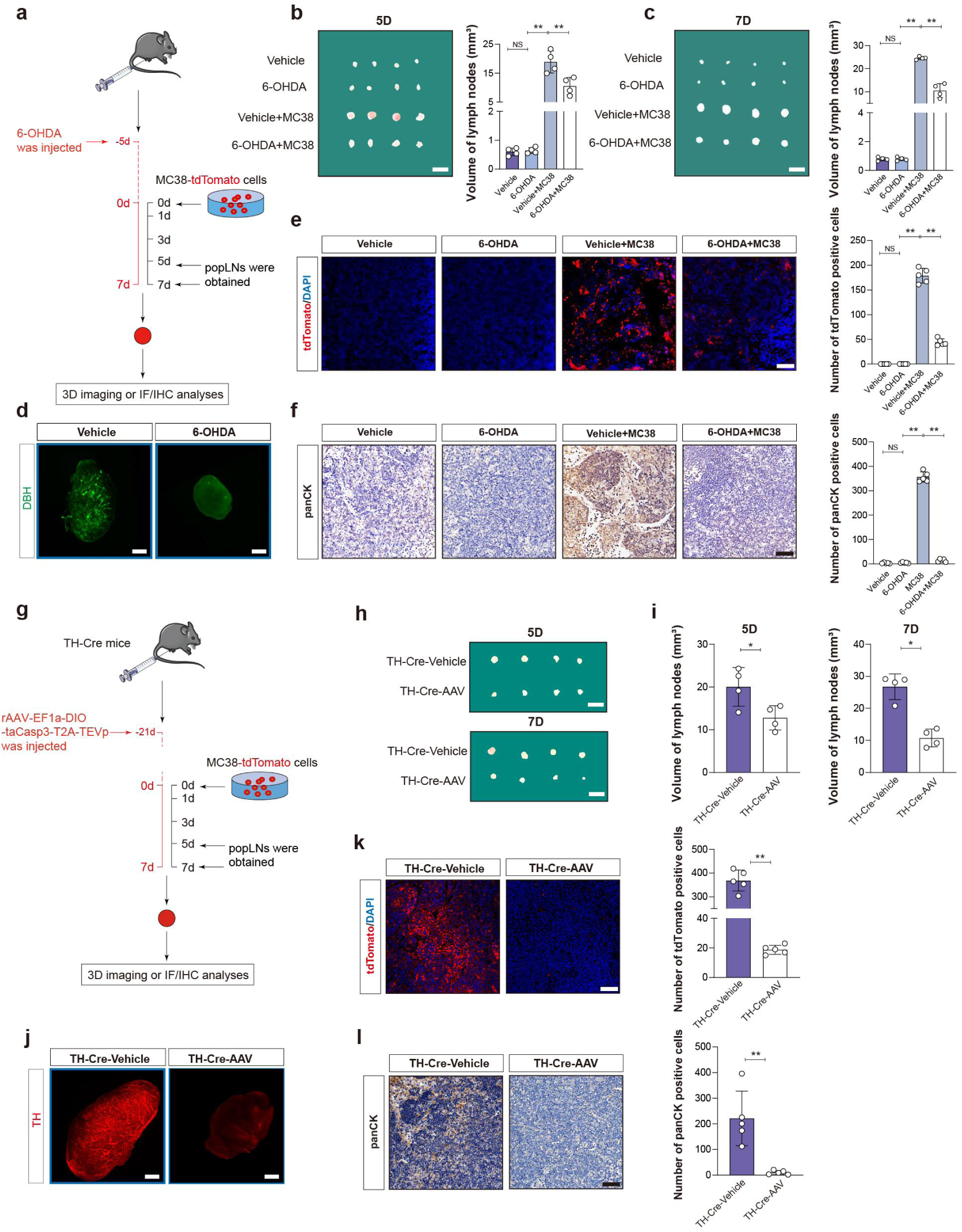
Local sympathetic innervation promotes tumor-induced LN remodeling and nodal metastasis. a, Schematic of local chemical sympathetic denervation. 6-OHDA or vehicle was administered to the fat pad surrounding the popLN five days before footpad inoculation of MC38-tdTomato cells. b, c, Representative images and quantification of popLN volume at days 5 (b) and 7 (c) after MC38-tdTomato inoculation in mice treated with local 6-OHDA or vehicle. Scale bars, 6 mm. d, Representative 3D images of DBH^+^ sympathetic nerve fibers in popLNs following local 6-OHDA-mediated denervation or vehicle treatment. Scale bar, 50 μm. e, Representative immunofluorescence images of tdTomato^+^ MC38 cells in popLNs following local 6-OHDA-mediated denervation or vehicle treatment, with corresponding quantification of tumor burden. Scale bar, 100 μm. f, Representative panCK immunohistochemical staining of the popLN medullary region following local 6-OHDA-mediated denervation or vehicle treatment, with corresponding quantification. Scale bar, 100 μm. g, Schematic of TH-Cre-dependent genetic sympathetic ablation. rAAV-EF1a-DIO-taCasp3-T2A-TEVp or control was locally delivered to the popliteal region of TH-Cre mice before footpad inoculation of MC38-tdTomato cells. h, i, Representative images (h) and quantification (i) of popLN volume at days 5 and 7 after MC38-tdTomato inoculation following TH-Cre-dependent sympathetic ablation or control treatment. Scale bars, 6 mm. j, Representative 3D images of TH^+^ sympathetic nerve fibers in popLNs following TH-Cre-dependent genetic sympathetic ablation or control treatment. Scale bar, 50 μm. k, Representative immunofluorescence images of tdTomato^+^ MC38 cells in popLNs following genetic sympathetic ablation, with corresponding quantification of tumor burden. Scale bar, 100 μm. l, Representative panCK immunohistochemical staining of the popLN medullary region following genetic sympathetic ablation, with corresponding quantification. Scale bar, 100 μm. **: *P*<0.01; NS: no significant difference.

To independently validate these findings using a genetic approach, we locally delivered rAAV-EF1a-DIO-taCasp3-T2A-TEVp to the popliteal region of TH-Cre mice to ablate sympathetic innervation before MC38 inoculation (**Fig. 7g**). Three-dimensional imaging confirmed a marked reduction of TH^+^ sympathetic fibers following genetic ablation (**Fig. 7j**). Consistent with the chemical denervation experiments, TH-Cre-dependent sympathetic ablation reduced popLN enlargement and nodal metastatic burden (**Fig. 7h, i, k, l**). Similar reductions in B220^+^ B cells, CD11b^+^ myeloid cells, and Ly6G^+^ neutrophils were observed (**Extended Data Fig. 12c, d**). Genetic sympathetic ablation was additionally associated with increased CD4^+^ and CD8^+^ T cells and reduced Treg abundance, whereas local 6-OHDA treatment produced more limited changes in T cell composition (**Extended Data Fig. 13a–l**). Importantly, neither local 6-OHDA-mediated sympathetic denervation nor TH-Cre-dependent genetic sympathetic ablation significantly altered primary footpad tumor burden at the examined time points (**Extended Data Fig. 14c, d**), arguing against reduced primary tumor growth as the explanation for the decreased nodal metastatic burden.

Collectively, these complementary chemical and genetic denervation experiments demonstrate that local sympathetic innervation functionally contributes to tumor-induced LN expansion, local immune remodeling, and nodal metastatic progression. Combined with the preceding stromal HGF and neuronal MET experiments, these findings support a mechanistic stromal HGF–MET–sympathetic axis linking stromal remodeling to neural plasticity and metastatic LN progression (**Fig. 8**).

**Fig. 8.**
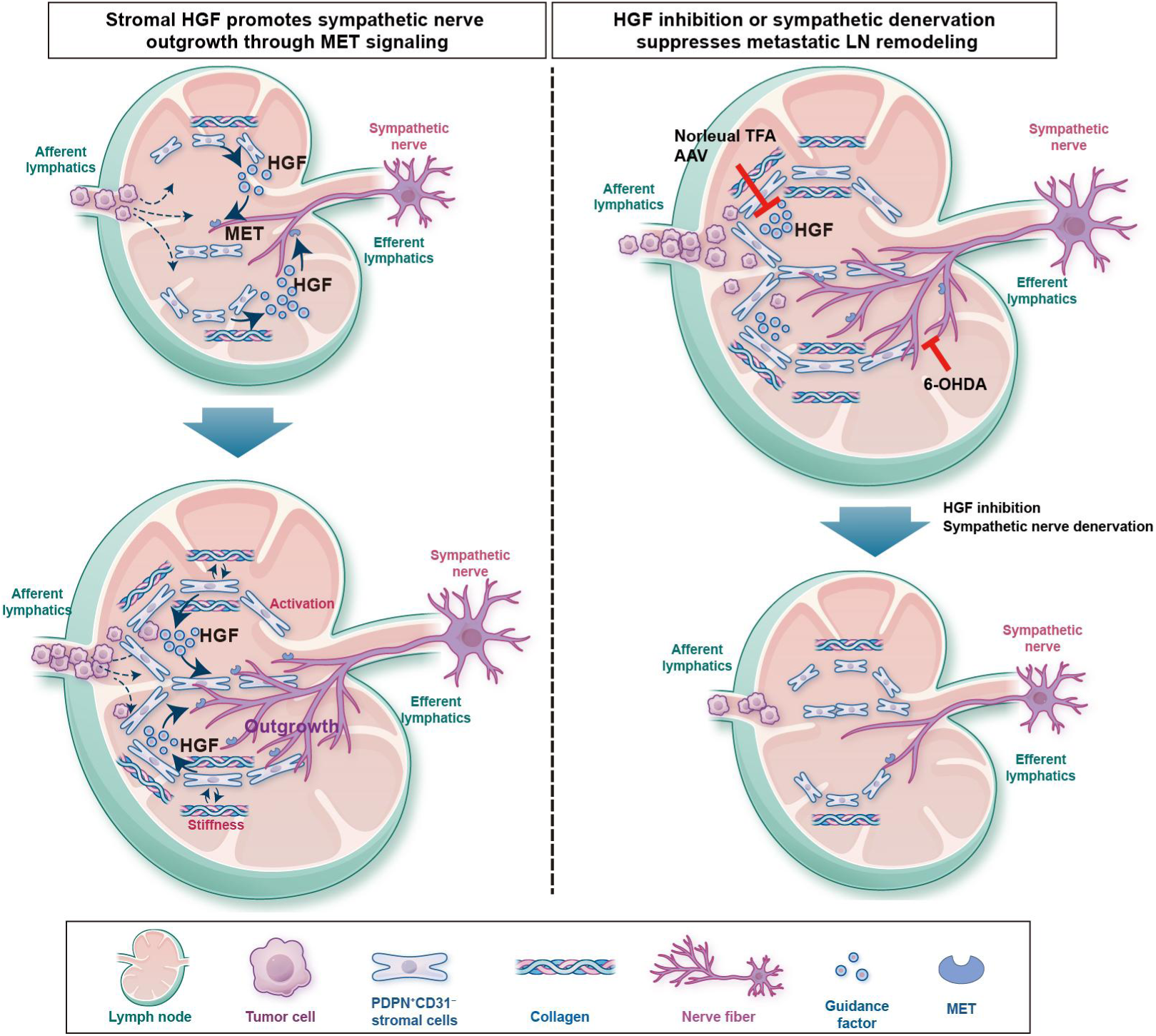
Proposed model of the stromal HGF–MET–sympathetic axis during metastatic LN remodeling. Afferent lymphatics enter the LN through the capsule, whereas efferent lymphatics exit at the hilum. Tumor-associated LN remodeling activates PDPN^+^ stromal cells, including FRCs, and is accompanied by progressive matrix stiffening. Stromal activation and increased matrix stiffness enhance HGF production, which signals through MET on sympathetic neurons to promote sympathetic nerve outgrowth. Sympathetic remodeling, in turn, contributes to LN expansion, local immune remodeling, and nodal metastatic progression. Local HGF inhibition or sympathetic denervation disrupts this stromal HGF–MET–sympathetic axis and reduces LN enlargement and nodal tumor progression.

## Discussion

Tumor-driven LN remodeling is accompanied by coordinated changes in local innervation. By integrating time-resolved whole-organ 3D imaging with single-nucleus transcriptomic and functional analyses, we resolved the temporal relationship between tumor infiltration, stromal remodeling and neural plasticity in TDLNs. Changes in the PDPN^+^ stromal network emerged early, followed by progressive tumor accumulation, LN expansion and increasing sympathetic nerve elongation and branching. Sensory innervation, by contrast, remained largely unchanged. This temporal pattern argues against increased innervation being simply a consequence of LN enlargement and instead identifies sympathetic remodeling as part of the evolving metastatic LN niche. Temporal snRNA-seq further linked these tissue-level changes to the emergence of neuro-supportive transcriptional programs in FRCs.

FRCs appear to be an important stromal source of signals that support this neural remodeling. During TDLN progression, FRCs acquired transcriptional programs associated with axon and neuronal projection development and increased HGF production. Local targeting of HGF in PDPN^+^ stromal cells suppressed sympathetic remodeling *in vivo*, whereas primary sympathetic neurons expressed MET and responded directly to HGF in a MET-dependent manner. These data support a stromal HGF–MET–sympathetic signaling axis in remodeling TDLNs. They also suggest a reciprocal relationship between lymphoid stroma and its innervation: neural inputs regulate lymphoid tissues, but pathological stromal remodeling can in turn reshape local neural architecture^2,4,18^.

Tissue mechanics provide an additional layer of regulation. Tumor-induced LN remodeling was accompanied by progressive collagen accumulation and matrix stiffening, consistent with the established role of FRC and ECM remodeling during LN expansion^9,10,12^. Increasing matrix stiffness experimentally enhanced FRC activation and HGF production. Mechanical remodeling may therefore be sensed by stromal cells and translated into trophic signals that promote neural growth. This provides a link between tissue mechanics and neural plasticity and places HGF within a broader stromal–mechanical–neural remodeling program.

The remodeled sympathetic innervation was also functionally relevant. Local chemical sympathectomy and TH-Cre-dependent genetic sympathetic ablation both reduced LN enlargement and nodal metastatic burden, indicating that sympathetic remodeling contributes to metastatic LN progression rather than simply accompanying it. Denervation also altered the immune composition of TDLNs. Previous studies have implicated sympathetic input in LN expansion, immune cell trafficking and retention, and local immune responses^2,7,8,22^. In our model, both denervation approaches altered myeloid and lymphoid populations, and genetic ablation was additionally associated with increased CD4^+^ and CD8^+^ T cells and reduced Treg abundance. How remodeled sympathetic fibers influence these populations remains unresolved and will require identification of the relevant neurotransmitters, receptors and cellular targets.

Our findings therefore extend the metastatic LN niche to include adaptive neural remodeling alongside its established immune and stromal components. The temporal framework developed here links tumor accumulation, stromal state, tissue mechanics and sympathetic innervation, while the HGF–MET axis provides a mechanism by which stromal remodeling can be coupled to neural growth. The upstream tumor-derived signals that initiate this stromal response, and the downstream adrenergic pathways through which remodeled nerves influence immunity and metastasis, remain to be defined. More generally, reciprocal adaptation between stroma and peripheral nerves may represent a feature of other tissues undergoing persistent pathological remodeling^24–26^.

## Data availability

All data are available upon request and relevant data are available in the Source Data. The snRNA-seq data from this study have been deposited in the Gene Expression Omnibus (GEO) with the accession code GSE280461.

## ACKNOWLEDGEMENTS

This study was supported by the National Natural Science Foundation of China (ID 82230087, 82350123, to Z.-G. Zhang; ID 82372589, 82073023, to J. Li; ID 82372821, 82103357 to L.-P. Hu; ID 82002485, to Q. Li), Key Laboratory of Systems Medicine for Cancer “Basic–Clinical Co-innovation Team” (SMC-BCC-01), the Shanghai Municipal Education Commission—Gaofeng Clinical Medicine Grant Support (ID 20181708, to Z.-G. Zhang), Innovative research team of high-level local universities in Shanghai (ID SHSMU-ZDCX20210802, to Z.-G. Zhang), Shanghai Pilot Program for Basic Research - Shanghai Jiao Tong University (ID 21TQ1400225, to Z.-G. Zhang), 111 project (ID B21024, to Z.-G. Zhang), Shanghai Eastern Talent Plan (ID QNWS2024104 to X.L. Zhang), Shanghai Science and Technology Commission Sailing Project (ID 22YF1445600 to D.-X. Li; ID 21YF1445200 to L.-P. Hu), the Shanghai Municipal Health Commission (ID 20214Y0200 to D.-X. Li), Key Areas Research and Development Programs of Guangdong Province (ID 2023B1111050009 to M.-J. Xu).

We thank Prof. Zhi-Feng Shao, Dr. Yan Yang, Dr. Xiao-Mei Yang, Dr. Yan-Li Zhang, Dr. Lei Zhu and Dr. Ni Zhang for assistance with our experiments.

## AUTHOR CONTRIBUTIONS

J.L. and Z.-G.Z. designed and supervised the overall study, analyzed data, and drafted the manuscript. J.L., X.Z., G.-H.S., L.-P.H., Q.L., R.L. and T.-H.W. constructed the mouse footpad model and collected the LN tissues. X.Z. and G.-H.S. performed the HYBRiD clearing and whole-mount tissue 3D imaging. J.L., H.L., Y.-Q.Z. and J.-X.X. analyzed the single-nucleus RNA sequencing data. X.Z., G.-H.S. and Y.-K.L. analyzed the data of 3D imaging. W.-J.H., Y.-Y.W., T.-S.B., F.W., S.Z. and C.-J.X. assisted with analysis of immunohistochemical and immunofluorescent staining. J.S., S.-Q.Y., X.-W.L. and J.X. technically assisted with experiments and analyzed data.

J.X. provided samples. J.L., S.-H.J., X.-L.Z., D.-X.L. and Z.-G.Z. supervised this study and edited the manuscript.

## Materials and Methods

### Cell culture

MC38 and PC12 cells, a catecholaminergic sympathetic neuron-like model, were purchased from the American Type Culture Collection (ATCC, USA) and cultured in DMEM (L110KJ, Basal Media, China) supplemented with 10% (v/v) fetal calf serum (DCF-201-0500, Dcell, China) and 1% penicillin/streptomycin (S110JV, Basal Media, China) at 37°C in a humidified incubator with 5% CO_2_. Serum-free solution for cryopreservation of cells (YC0100, Yoche Biotechnology, Shanghai, China) was used in this experiment.

### Culture Medium

Primary FRCs or MC38 cells were seeded in 10-cm dishes and cultured in complete medium until reaching the indicated confluence. The medium was then replaced with serum-free medium, and cells were further incubated for 48 h. The conditioned medium was subsequently collected, centrifuged to remove cellular debris, and stored at −20°C until use.

### Antibodies

Primary and secondary antibodies were used according to the manufacturers’ instructions. Detailed information on all antibodies used for whole-mount immunofluorescence staining, tissue and cell immunofluorescence staining, immunohistochemistry and flow cytometry, including target, fluorophore, host species, clone, manufacturer, catalogue number, application and working dilution or concentration, is provided in **Extended Data Table 1**.

### Real-time qPCR

Real-time qPCR reactions were prepared using 2 × Universal Blue SYBR Green qPCR Master Mix (G3326, Servicebio, China) and performed on a 7500 Real-Time PCR System (Applied Biosystems, USA). Thermal cycling consisted of an initial denaturation at 95 °C for 10 min, followed by 40 cycles of 95 °C for 15 s and 60 °C for 30 s. Relative mRNA expression was calculated using the 2^−ΔΔCt^ method and normalized to 18S mRNA levels. Primer sequences are listed below:

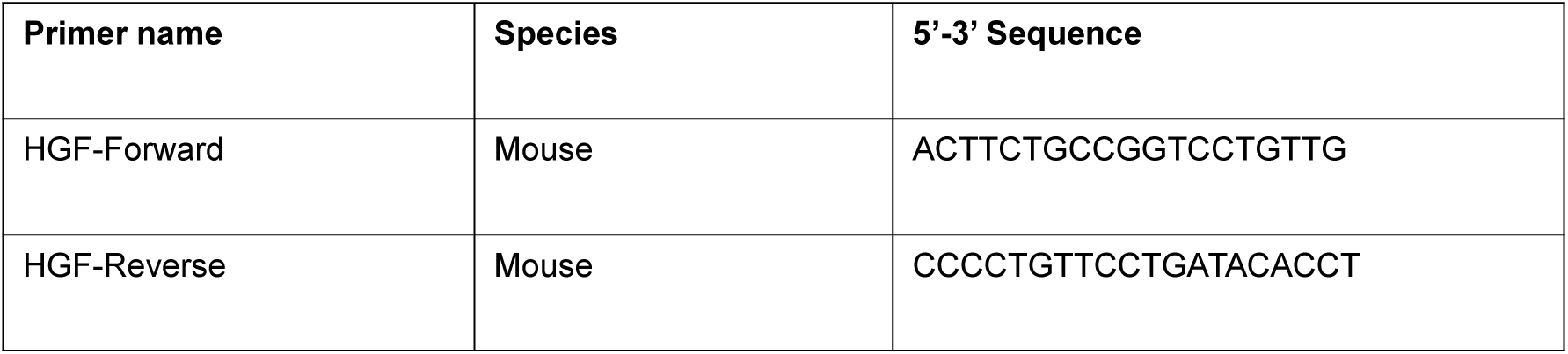
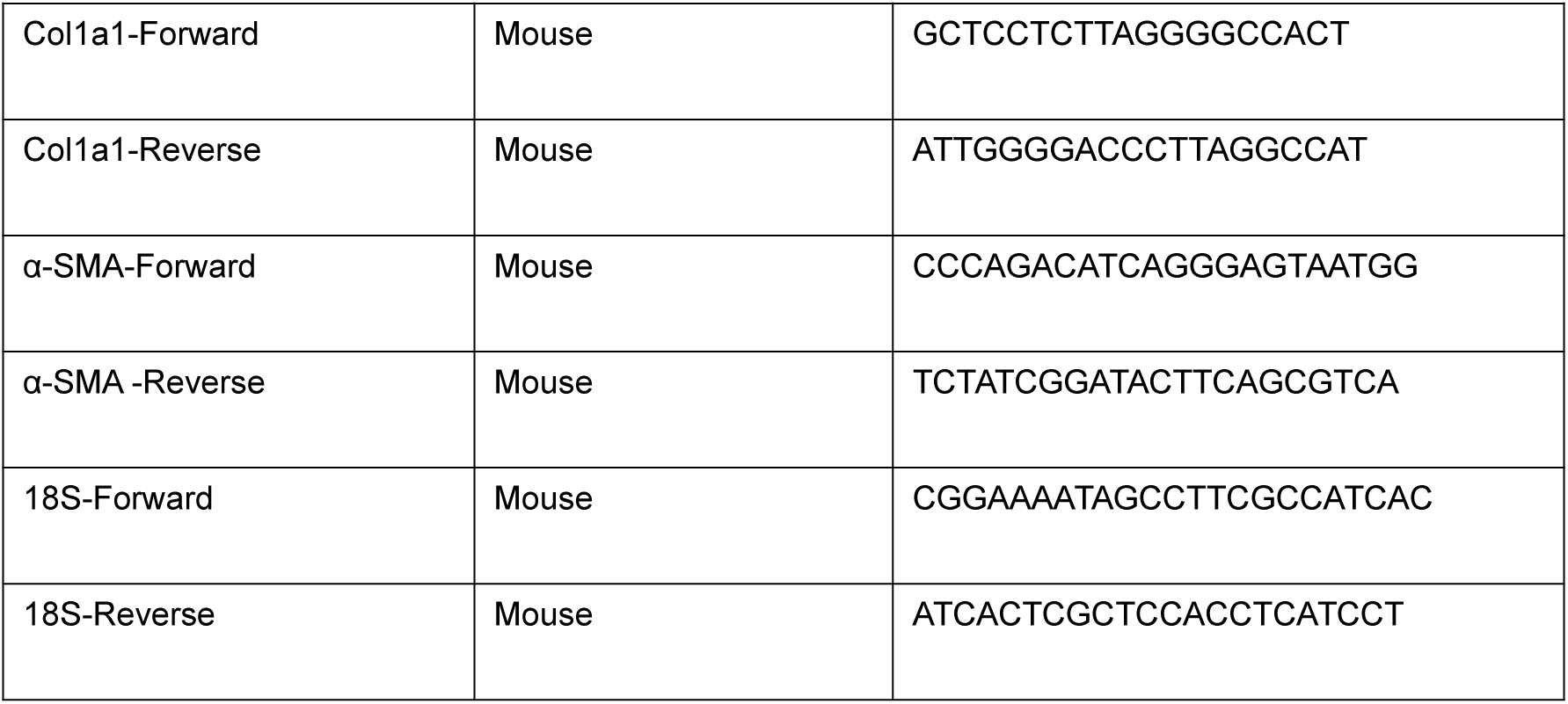

### Enzyme-Linked Immunosorbent Assay (ELISA)

The required ELISA plate (PCDBM0133, PC-biotech, Shanghai, China) was taken out and equilibrated to room temperature. 50 μl each standard samples of different concentrations were added into the standard wells in sequence. 50 μl test samples were added into the other wells. After incubation at 37℃ for 90 min, each well was filled with 350 μl washing solution and stood for 1 min. The washing solution was then discarded, and the plate was blotted dry on absorbent paper. This washing procedure was repeated three times.

Then 100 μl biotin-labeled detection antibody was added into each well. The reaction wells were covered with a sealing film. After incubation at 37℃ for 60 min, the liquid was discarded. Each well was washed with 350 μl washing solution 3 times.

Subsequently, 100 μl of horseradish peroxidase (HRP)-conjugated streptavidin, diluted according to the manufacturer’s instructions, was added to each well. The plate was covered with a sealing film and incubated at 37 °C for 30 min. The solution was then discarded, and each well was washed five times with 350 μL of washing buffer.

Finally, 50 μl TMB solution was added into each well. After incubation at 37℃ in the dark for 15 min, the sealing film was removed and 50 μL stop solution was added to each well. Optical density (OD) was measured at 450 nm. A four-parameter logistic curve was fitted to the standards, and sample concentrations were calculated from the corresponding OD450 values.

### Animals

Male C57BL/6J mice (6–8 weeks old) were purchased from GemPharmatech Co., Ltd. (Nanjing, China). *Hgf ^fl/fl^* mice on a C57BL/6JGpt background were generated by GemPharmatech Co., Ltd. using a Cas9-based conditional knockout strategy, in which loxP sites were introduced to flank exon 3 of *Hgf*, allowing Cre-dependent deletion of the targeted region. TH-Cre, Dbh-P2A-EGFP, and Calca-P2A-BFP mice were also obtained from GemPharmatech Co., Ltd. The Dbh-P2A-EGFP^1^ and Calca-P2A-BFP^2^ reporter mouse lines were generated using CRISPR/Cas9-mediated knock-in strategies^3^. In the Calca-P2A-BFP line, a P2A-BFP^4^ cassette was inserted in frame immediately upstream of the endogenous TAA stop codon in exon 4. The guide RNA sequences were 5′-AGTTGCCAAAATGGGATTGG-3′ and 5′-AGGAGCTTAGTTGCCAAAAT-3′. In the Dbh-P2A-EGFP line, a P2A-EGFP^4^ cassette was inserted in frame immediately upstream of the endogenous TAG stop codon in exon 12. The guide RNA sequences were 5′-CTCTGTTGTGATGGGCACAG-3′ and 5′-CTGTTGTGATGGGCACAGTG-3′.

The corresponding endogenous 3′ untranslated region sequences were retained in both targeting designs.

The donor plasmids were GJS022022065851-01-Calca-P2A-BFP (8,026 bp) and GJS072021032297C2-01-Dbh-P2A-EGFP (8,140 bp). The Calca donor plasmid contained an MC1 promoter-driven diphtheria toxin A (DTA) cassette followed by a rabbit β-globin polyadenylation signal. This cassette was present in the donor plasmid but was not included in the designed targeted genomic sequence. No DTA cassette was annotated in the Dbh donor plasmid map. Donor plasmids were validated by PCR and restriction enzyme digestion. Founder mice were identified by PCR and sequencing and subsequently maintained for experimental use. Mice were housed under specific pathogen-free conditions and handled in accordance with protocols approved by the Laboratory Animal Center of Shanghai Jiao Tong University. All animal experiments were approved by the Laboratory Animal Center of Shanghai Jiao Tong University (approval no. A2024402). All animals received humane care according to the Guide for the Care and Use of Laboratory Animals published by the National Institutes of Health (NIH).

### Mouse footpad model

A popliteal lymph node (popLN) metastasis model was established by injecting MC38 or MC38-tdTomato cells into the footpad of C57BL/6J mice. Briefly, 4×10^6^ cells suspended in 25 μL PBS were injected into the footpad of each mouse. popLNs were harvested at the indicated time points (day 0, 1, 2, 3, 5, 7, and 10) after tumor inoculation for subsequent analyses. For fluorescence-based tracing experiments, MC38-tdTomato cells were used. Mice were randomly assigned to experimental groups.

### Local HGF-targeting interventions

For local inhibition of HGF signaling, Norleual TFA was injected into the popliteal fat pad at a concentration of 0.1 mg/mL in a volume of 50 μL per lymph node 6 h before MC38 cell inoculation. Additional injections were administered every 3 days thereafter. Vehicle-treated mice were used as controls.

For local *Hgf* silencing, an AAV9 vector carrying an Hgf-targeting shRNA (AAV-sh-*Hgf*; order no. V3000970-1) and the corresponding empty-vector control were obtained from Shanghai Share-Bio Technology Co., Ltd. (Shanghai, China). Both vectors used the VA060-AAV-U6-MCS-CMV-ZsGreen backbone. The *Hgf*-targeting shRNA was expressed under the U6 promoter, whereas ZsGreen expression was driven by the CMV promoter. The *Hgf*-targeting sequence was 5′-CTTCGAGCTATCGCGGTAAAG-3′. The shRNA insert was cloned between the EcoRI and BamHI sites and verified by Sanger sequencing. AAV-sh-*Hgf* or the corresponding empty-vector control was injected into the popliteal fat pad at a titer of 8.5 × 10¹² vg/mL in a volume of 50 μL per popLN corresponding to 4.25 × 10¹¹ vg per popLN, 21 days before MC38 cell inoculation. Popliteal lymph nodes were harvested at the indicated time points for subsequent analyses.

For local targeting of *Hgf* in PDPN^+^ stromal cells, AAV9-PDPN-Cre was injected into the popliteal fat pad of *Hgf* ^fl/fl^ mice at a titer of 8.6 × 10^12^ vg/mL in a volume of 50 μL per lymph node 21 days before MC38 cell inoculation. Vehicle mice received the corresponding control treatment. popLNs were harvested at the indicated time points for subsequent analyses.

### Chemical sympathetic denervation

For local sympathetic denervation, 6-hydroxydopamine (6-OHDA) (HY-B1081A, China, MedChemExpress) was prepared at 20 mg/mL in saline containing 0.1% ascorbic acid. A total volume of 50 μL was injected into the popliteal fat pad adjacent to each popLN on each side. The first injection was administered 5 days before MC38 cell inoculation, followed by additional injections every 3 days thereafter^5^. 6-OHDA was dissolved in saline containing 0.1% ascorbic acid to prevent oxidation. Control mice received vehicle only^6,7^. popLNs were collected at the indicated time points for further analysis.

### Genetic sympathetic ablation

For genetic ablation of sympathetic nerves, TH-Cre mice were locally injected in the popliteal fat pad with AAV9-EF1α-DIO-taCasp3-T2A-TEVP (8.0 × 10^12^ vg/mL, 50 μL per lymph node) 21 days before MC38 cell inoculation. Control mice received the corresponding control treatment. popLNs were harvested at the indicated time points for subsequent analyses, including lymph node size, whole-mount 3D imaging, and evaluation of metastatic burden.

### Sample pretreatment

At the indicated time points, mice were euthanized and popLNs were carefully dissected for downstream analyses. For whole-mount staining and tissue clearing, lymph nodes were fixed overnight in 4% formaldehyde at 4°C. After fixation, samples were washed in PBS and stored in 1 × PBS containing 0.02% NaN3 until further processing. At least five mice were used at each time point.

### Preparation of tissue-clearing reagents

In a fume hood, N, N, N’, N’-Tetrakis (2-hydroxypropyl) ethylenediamine (122262, Sigma, USA) was diluted to a final concentration of 25% as a decoloring buffer with 10 × PBS (70011044, Thermo Fisher Scientific, USA) and distilled water and stirred with a magnetic stirrer. The mixture was stored at room temperature in a dark place. THF (T103264, Aladdin, China) gradients of 50%, 70%, 80%, and 95% were prepared using the decoloring buffer and stored at 4 °C. 1 × PBST wash buffer containing 0.1% Tween-20 (T8220, Solarbio, China) and 0.02% NaN3 was prepared. 1 × PBST staining buffer containing 0.2% Triton X-100 (T8200, Solarbio, China) was prepared.

### Tissue clearing

Initially, the samples were washed with 1 × PBS 3 times to remove any residual formaldehyde, with each washing session lasting a minimum of 1 hour. Samples were incubated in decolorizing buffer at 37 °C with gentle agitation for 2 days. The decolorizing solution was changed once a day. Decolorization was extended when necessary.

Following the decolorizing phase, the samples underwent an additional three rounds of thorough washing with 1 × PBS, each session lasting one hour. Samples were sequentially incubated in 50%, 70%, 80%, and 95% THF at 4 °C with agitation for at least 30 min to perform tissue clearing. The samples were then put into a 100% dichloromethane (270997, Sigma, USA) solution for tissue clearing and shaken for 60 min at 4 °C. The dichloromethane solution was replaced once.

The samples were then placed in 95% THF, 80% THF, 70% THF, and 50% THF in order and then put on a 4°C shaker for at least 30 min for each gradient to perform tissue clearing. The samples were washed with 1 × PBS three times for one hour each time. Before imaging, the samples were put in the Easy Index (RI=1.52, CAT#: EI-500-1.52, Lifecanvas Technologies, USA) and shaken overnight at 37 °C. The next day, the samples were placed on a shaker at room temperature for refractive index matching^8^.

### Sample staining

For whole-mount immunofluorescence staining, cleared lymph nodes were incubated overnight in 5% bovine serum albumin (BSA) in PBS at 4°C with gentle agitation. Samples were then washed three times with PBS for 1 h per wash at room temperature. Primary antibodies against TH, PGP9.5 and NFL were incubated with the samples in a total volume of 200 μL for 7 days at 37°C with gentle agitation. Samples were subsequently washed three times with 1 × PBS for 1 h per wash and incubated with the corresponding secondary antibodies for 7 days at 37°C with gentle agitation.

### Atomic force microscopy (AFM)

The mechanical stiffness of popLNs was measured by atomic force microscopy using a MFP-3D-BIO AFM (Asylum Research, Oxford Instruments, UK) equipped with a spherical probe (5 μm bead radius, spring constant ∼0.06 N/m). Fresh lymph nodes were embedded in low-melting agarose, sectioned into 200-μm slices, and measured in PBS at room temperature. Force–indentation curves were acquired at an approach speed of 2 μm/s with a trigger force of 2 nN, resulting in an indentation depth of approximately 2 μm. For each lymph node, measurements were collected from at least 5 independent regions, with 5–10 force curves per region. Young’s modulus was calculated by fitting the force–indentation curves using the Hertz contact model with a Poisson’s ratio of 0.5.

### Light sheet fluorescence microscopy (LSFM) imaging

Cleared lymph nodes were embedded in 1–2% low-melting-point agarose prepared in EasyIndex and mounted for light-sheet fluorescence microscopy. Samples were imaged using a MegaSPIM light-sheet microscope (LifeCanvas Technologies, USA) equipped with a 9 × objective, with a voxel size of 0.72 μm. Laser power and other acquisition settings were kept constant across samples within each experiment.

### Image processing and quantization

Three-dimensional image reconstruction and quantitative analysis were performed using Imaris 10.1 software (Oxford Instruments, UK)^9^. Positive signals in each fluorescence channel were segmented using the Surface module, and nerve fibers were reconstructed using the Filament module. Quantification of nerve remodeling was based on Filament Length (sum) and Filament No. Branch Points. PDPN-positive stromal networks were quantified as positive area using the Surface module. tdTomato-positive metastatic tumor cells were quantified based on the number of positive cells, as indicated in each experiment. The same analysis workflow and thresholding criteria were applied to all samples within each dataset.

### Immunofluorescence (IF) staining

For cell immunofluorescence staining, 2 × 10³ cells per well were seeded in 24-well plates. Cells were fixed with 4% paraformaldehyde (PFA), blocked with 5% bovine serum albumin (BSA) in PBS, and incubated with primary antibodies against TH, MET as indicated, followed by the appropriate fluorophore-conjugated secondary antibodies. DAPI (C0060, Solarbio, China) was used for nuclear counterstaining for 10 min at room temperature.

For tissue immunofluorescence staining, paraffin-embedded sections were deparaffinized and rehydrated through a graded ethanol series, followed by antigen retrieval in citrate-based buffer. Sections were blocked with 5% BSA for 1 h and incubated with primary antibodies against TH, NFL, PGP9.5, MET, PDPN or HGF, as indicated, at 4°C overnight. Sections were subsequently incubated with the appropriate fluorophore-conjugated secondary antibodies for 1 h at room temperature. DAPI was used to counterstain nuclei for 5 min. Images were acquired using a Leica confocal microscope (Leica, Germany). Detailed information on the antibodies and working dilutions is provided in **Extended Data Table 1**.

### Immunohistochemical staining

Paraffin-embedded lymph node sections were dewaxed in xylene and rehydrated through a graded ethanol series. Antigen retrieval was performed in citrate-based buffer. After blocking with 5% bovine serum albumin (BSA), sections were incubated with primary antibodies against panCK and α-SMA at 4°C overnight, followed by incubation with the appropriate HRP-conjugated secondary antibodies. Immunoreactivity was visualized using a DAB chromogenic system, and nuclei were counterstained with hematoxylin. Images were acquired under a light microscope. Quantification was performed based on the positive staining area or signal intensity, as indicated in each experiment.

### Single-nucleus RNA sequencing (snRNA-seq)

At each time point, 10 popLNs collected from five mice were pooled as one sample for snRNA-seq to meet the input requirements for nuclei isolation and library construction. snRNA-seq was performed according to the experimental procedure of Shanghai Bohao Biotechnology Co., Ltd. Libraries were generated using the 10x Genomics Chromium Single Cell 3′ v3 chemistry. Raw sequencing data were processed using Cell Ranger v7.0.1 (10 × Genomics) with intronic reads included and aligned to the mm10-1.2.0 mouse reference transcriptome. Data were subsequently preprocessed and normalized before downstream clustering and cell-type annotation. After integration of multiple samples, the Louvain algorithm was used for clustering, and t-SNE was used for visualization of the high-dimensional data in two dimensions. Cell types were initially annotated using SingleR and were further validated based on canonical marker genes. SingleR-based annotations were subsequently refined according to the expression of established cell-type-specific marker genes reported in previous studies. Detailed classification of individual cell populations was performed based on canonical marker genes and previous studies.^10,11^.

The snRNA-seq data from this study have been deposited in the Gene Expression Omnibus (GEO) with the accession code GSE280461.

### Flow cytometry

popLNs were harvested at the indicated time points after MC38 cell inoculation and mechanically dissociated into single-cell suspensions. Samples were digested at 37°C for 30 min in DMEM containing 10% FBS, 1% penicillin/streptomycin, Collagenase type I (C5894, Sigma-Aldrich, Germany; 0.05 mg/mL), Collagenase type IV (C1889, Sigma-Aldrich, Germany; 0.05 mg/mL), Hyaluronidase (H4272, Sigma-Aldrich, Germany; 0.025 mg/mL), and DNase I (11284932001, Roche, Switzerland; 0.01 mg/mL). Digested samples were filtered through a 70 μm cell strainer and stained with fluorochrome-conjugated antibodies. Dead cells were excluded using a viability dye. Data were acquired on an LSRFortessa flow cytometer (BD Biosciences, USA) and analyzed using FlowJo software (Tree Star, USA).

For flow cytometric analysis, the following antibodies and viability dye were used according to the indicated panels: PE anti-mouse CD45, FITC anti-mouse CD4, Brilliant Violet 510 anti-mouse CD8a, APC anti-mouse FOXP3, PE/Cyanine7 anti-mouse NK-1.1, and Zombie NIR Fixable Viability Kit (423106, BioLegend, USA) for panel 1; PerCP anti-mouse CD45, Brilliant Violet 421 anti-mouse/human CD45R/B220, PE anti-mouse CD11c, PE/Cyanine7 anti-mouse I-A/I-E, and Zombie NIR Fixable Viability Kit (423106, BioLegend, USA) for panel 2; PerCP anti-mouse CD45, BUV805 anti-mouse CD11b, FITC anti-mouse Ly-6C, PE anti-mouse Ly-6G, BUV661 anti-mouse F4/80, and Zombie NIR Fixable Viability Kit (423106, BioLegend, USA) for panel 3; and PerCP anti-mouse CD45, APC anti-mouse CD31, and PE anti-mouse Podoplanin for panel 4.

Cells were first gated on FSC/SSC to exclude debris, followed by singlet gating and live-cell gating. CD45 was used to distinguish immune cells from stromal cells. PDPN⁺CD31⁻ stromal cells were gated from the CD45⁻ fraction. Among CD45⁺ immune cells, B cells were identified as B220⁺, CD4⁺ and CD8⁺ lymphocyte populations were identified based on CD4 and CD8 expression within the lymphocyte gate, NK cells as NK1.1⁺, neutrophils as CD11b⁺Ly6G⁺, monocytes as CD11b⁺Ly6C⁺, macrophages as CD11b⁺F4/80⁺, and dendritic cells (DCs) as CD11c⁺MHC II⁺. The relative proportion of each population was calculated as indicated in each experiment. Cell numbers were normalized to 1 × 10^4^ viable cells from each popLN and the corresponding flow-cytometric frequency of each cell population. Relative frequencies and cell numbers were reported as indicated in each experiment.

### Primary FRC isolation and culture

Draining lymph nodes (LNs) were harvested from ten 4-week-old mice and washed twice with sterile PBS prechilled to 4 ℃. The LNs were then finely minced into approximately 1 mm ³ fragments using sterile scissors. Tissue fragments were digested in 1 mL of 0.2 mg/mL Collagenase P (11213865001, Roche, USA) at 37℃ for 50 min, with gentle agitation every 10 min. Following digestion, the tissue suspension was gently triturated with a 1-mL pipette tip for 1 min. Larger undigested fragments were allowed to settle for 1 min, after which the supernatant was collected.

The collected supernatant was diluted 1:1 with prechilled DMEM/F-12 (1:1) (L310KJ, BasalMedia, China) supplemented with 10% fetal bovine serum (FBS) and 1% antibiotic solution (S110JV, BasalMedia, China). The suspension was centrifuged at 300×g for 5 min at 4℃, and the resulting cell pellet was resuspended in complete DMEM/F-12 medium. Cells were seeded into 25-cm² culture flasks and maintained at 37 ℃ in a humidified atmosphere containing 5% CO2. The culture medium was replaced after 12 h and subsequently every 3 days. Before each medium change, adherent cells were gently washed with DMEM/F-12 (1:1).

After approximately 1 week of culture, adherent cells were washed with calcium- and magnesium-free PBS and detached using 0.05% trypsin (S320KJ, BasalMedia, China) for 120 s. The detached cells were transferred to fresh 25-cm² culture flasks and maintained under the same culture conditions. This passaging procedure was repeated for four cycles.

### Primary sympathetic neuron culture and MET knockdown

Primary sympathetic ganglia were isolated from 1-week-old DBH-P2A-EGFP reporter mice or C57BL/6J mice, dissociated using Collagenase P (11213865001, Roche, Switzerland) at 0.2 mg/mL, and cultured on plates coated with Poly-D-Lysine (A3890401, Gibco, USA) and Laminin, Mouse Protein, Natural (23017015, Gibco, USA) in Neurobasal Medium (21103049, Gibco, USA) supplemented with B-27 Supplement (17504044, Gibco, USA; 1X), GlutaMAX Supplement (35050061, Gibco, USA; 1X), and Mouse NGF Recombinant Protein (A42627, Invitrogen, USA; 50 ng/mL). For MET knockdown, primary sympathetic neurons cultured in 24-well plates were transfected with 50 pmol MET-targeting siRNA or the corresponding negative control using the ProteanFect Max Transfection Kit (PT02, Nano Portal Biotech, China) according to the manufacturer’s instructions for primary cells with minor modifications. Briefly, 50 pmol siRNA was mixed with 200 μL Reagent A (PT02), followed by addition of 3.5 μL Reagent B (PT02) and 40 μL Reagent C (PT02). Cells at 50%–80% confluence were washed twice with serum-free DMEM and maintained in 100 μL serum-free DMEM during transfection. The transfection mixture was added directly to the cells and incubated at 37°C for 30 min, after which the mixture was removed and replaced with Neurobasal complete medium. Cells were used for subsequent assays 48 h after transfection. The MET-targeting construct was sequence-verified before use. Recombinant mouse HGF (315-23-20UG, PeproTech, USA) was added at 50 ng/mL for 24 h^12^, followed by immunofluorescence staining and neurite outgrowth analysis. The sequences of the mouse MET-targeting siRNA were as follows: Met-mouse-1847, sense 5′-GCACGACAAAUACGUUGAATT-3′ and antisense 5′-UUCAACGUAUUUGUCGUGCTT-3′. The negative control siRNA sequences were sense 5′-UUCUCCGAACGUGUCACGUTT-3′ and antisense 5′-ACGUGACACGUUCGGAGAATT-3′.

### Statistical analysis

Data are presented as the means ± standard errors of the mean (SEMs). Statistical analyses were performed using GraphPad Prism (version 8.0.1.244 x64, USA) for Windows. Comparisons between groups were performed using two-tailed Student’s t-tests or one-way ANOVA followed by Dunnett, as specified in the corresponding figure legends. Pearson correlation analysis was used for correlation analyses. Fisher’s exact test was used to assess associations between categorical variables in Fig. 2e, f. P < 0.05 was considered statistically significant.

### Inclusion and ethics

 This study involved animal subjects and was approved by the Laboratory Animal Center of Shanghai Jiao Tong University (ID: A2024402).

## References

1. Reticker-Flynn, N. E. & Engleman, E. G. Lymph nodes: at the intersection of cancer treatment and progression. Trends in Cell Biology 33, 1021–1034 (2023).

2. De Virgiliis, F., Oliva, V. M., Kizil, B. & Scheiermann, C. Control of lymph node activity by direct local innervation. Trends in Neurosciences 45, 704–712 (2022).

3. Krishnamurty, A. T. & Turley, S. J. Lymph node stromal cells: cartographers of the immune system. Nat Immunol 21, 369–380 (2020).

4. Huang et al. Lymph nodes are innervated by a unique population of sensory neurons with immunomodulatory potential. Cell 184, 441–459.e25 (2021).

5. Cleypool, C. G. J., Mackaaij, C., Lotgerink Bruinenberg, D., Schurink, B. & Bleys, R. L. A. W. Sympathetic nerve distribution in human lymph nodes. Journal of Anatomy 239, 282–289 (2021).

6. Hanč, P. et al. Multimodal control of dendritic cell functions by nociceptors. Science 379, eabm5658 (2023).

7. Chen, C.-S. et al. Loss of direct adrenergic innervation after peripheral nerve injury causes lymph node expansion through IFN-γ. Journal of Experimental Medicine 218, e20202377 (2021).

8. Klatt, S. et al. Peripheral elimination of the sympathetic nervous system stimulates immunocyte retention in lymph nodes and ameliorates collagen type II arthritis. Brain, Behavior, and Immunity 54, 201–210 (2016).

9. Acton, S. E. et al. Dendritic cells control fibroblastic reticular network tension and lymph node expansion. Nature 514, 498–502 (2014).

10. Astarita, J. L. et al. The CLEC-2–podoplanin axis controls the contractility of fibroblastic reticular cells and lymph node microarchitecture. Nat Immunol 16, 75–84 (2015).

11. De Martin, A., Stanossek, Y., Pikor, N. B. & Ludewig, B. Protective fibroblastic niches in secondary lymphoid organs. J Exp Med 221, e20221220 (2024).

12. Riedel, A., Shorthouse, D., Haas, L., Hall, B. A. & Shields, J. Tumor-induced stromal reprogramming drives lymph node transformation. Nat Immunol 17, 1118–1127 (2016).

13. Du Bois, H., Heim, T. A. & Lund, A. W. Tumor-draining lymph nodes: At the crossroads of metastasis and immunity. Sci. Immunol. 6, eabg3551 (2021).

14. Nudell, V. et al. HYBRiD: hydrogel-reinforced DISCO for clearing mammalian bodies. Nat Methods 19, 479–485 (2022).

15. Li, J. et al. Spatiotemporal quantification of metastatic tumour cell growth and distribution in lymph nodes by whole-mount tissue 3D imaging. Int J Biol Sci 18, 3993–4005 (2022).

16. Rodda et al. Single-Cell RNA Sequencing of Lymph Node Stromal Cells Reveals Niche-Associated Heterogeneity. Immunity 48, 1014–1028.e6 (2018).

17. Kapoor, V. N. et al. Gremlin 1+ fibroblastic niche maintains dendritic cell homeostasis in lymphoid tissues. Nat Immunol 22, 571–585 (2021).

18. Cheong, L. Y. et al. Fibroblastic reticular cells in lymph node potentiate white adipose tissue beiging through neuro-immune crosstalk in male mice. Nat Commun 14, 1213 (2023).

19. David, M. D. et al. Signalling by neurotrophins and hepatocyte growth factor regulates axon morphogenesis by differential β-catenin phosphorylation. Journal of Cell Science 121, 2718–2730 (2008).

20. Hartmann, S., Bhola, N. E. & Grandis, J. R. HGF/Met Signaling in Head and Neck Cancer: Impact on the Tumor Microenvironment. Clinical Cancer Research 22, 4005–4013 (2016).

21. Renz, B. W. et al. β2 Adrenergic-Neurotrophin Feedforward Loop Promotes Pancreatic Cancer. Cancer Cell 34, 863–867 (2018).

22. Reel, J. M. et al. The Sympathetic Nervous System Is Necessary for Development of CD4+ T-Cell Memory Following *Staphylococcus aureus* Infection. The Journal of Infectious Diseases 228, 966–974 (2023).

23. Li, Y. et al. An HGF-dependent positive feedback loop between bladder cancer cells and fibroblasts mediates lymphangiogenesis and lymphatic metastasis. Cancer Commun 43, 1289–1311 (2023).

24. Wang, Y. et al. The role of somatosensory innervation of adipose tissues. Nature 609, 569–574 (2022).

25. Zhang, Y. et al. Cancer cells co-opt nociceptive nerves to thrive in nutrient-poor environments and upon nutrient-starvation therapies. Cell Metabolism 34, 1999–2017.e10 (2022).

26. Banh, R. S. et al. Neurons Release Serine to Support mRNA Translation in Pancreatic Cancer. Cell 183, 1202–1218.e25 (2020).

## References

1. Torres, H. et al. Sympathetic innervation of the mouse kidney and liver arising from prevertebral ganglia. *American Journal of Physiology-Regulatory*, Integrative and Comparative Physiology 321, R328–R337 (2021).

2. McCoy, E. S., Taylor-Blake, B. & Zylka, M. J. CGRPα-Expressing Sensory Neurons Respond to Stimuli that Evoke Sensations of Pain and Itch. PLoS ONE 7, e36355 (2012).

3. Yang, H. et al. One-step generation of mice carrying reporter and conditional alleles by CRISPR/Cas-mediated genome engineering. Cell 154, 1370–1379 (2013).

4. Trichas, G., Begbie, J. & Srinivas, S. Use of the viral 2A peptide for bicistronic expression in transgenic mice. BMC Biol 6, 40 (2008).

5. Klatt, S. et al. Peripheral elimination of the sympathetic nervous system stimulates immunocyte retention in lymph nodes and ameliorates collagen type II arthritis. Brain, Behavior, and Immunity 54, 201–210 (2016).

6. Blank, C. L. et al. Intracyclization rates of 6-hydroxydopamine and 6-aminodopamine analogs under physiological conditions. J Med Chem 19, 178–180 (1976).

7. Wu, X.-H. et al. The Role of Retinal Dopamine in C57BL/6 Mouse Refractive Development as Revealed by Intravitreal Administration of 6-Hydroxydopamine. Invest Ophthalmol Vis Sci 57, 5393–5404 (2016).

8. Nudell, V. et al. HYBRiD: hydrogel-reinforced DISCO for clearing mammalian bodies. Nat Methods 19, 479–485 (2022).

9. Li, J. et al. Spatiotemporal quantification of metastatic tumour cell growth and distribution in lymph nodes by whole-mount tissue 3D imaging. Int J Biol Sci 18, 3993–4005 (2022).

10. Rodda, L. B. et al. Single-Cell RNA Sequencing of Lymph Node Stromal Cells Reveals Niche-Associated Heterogeneity. Immunity 48, 1014–1028.e6 (2018).

11. Kapoor, V. N. et al. Gremlin 1+ fibroblastic niche maintains dendritic cell homeostasis in lymphoid tissues. Nat Immunol 22, 571–585 (2021).

12. David, M. D. et al. Signalling by neurotrophins and hepatocyte growth factor regulates axon morphogenesis by differential β-catenin phosphorylation. Journal of Cell Science 121, 2718–2730 (2008).

