## Supplemental Figures for "Stromal control of sympathetic rewiring in metastatic lymph nodes"

**Extended Data Figures and figure legends**

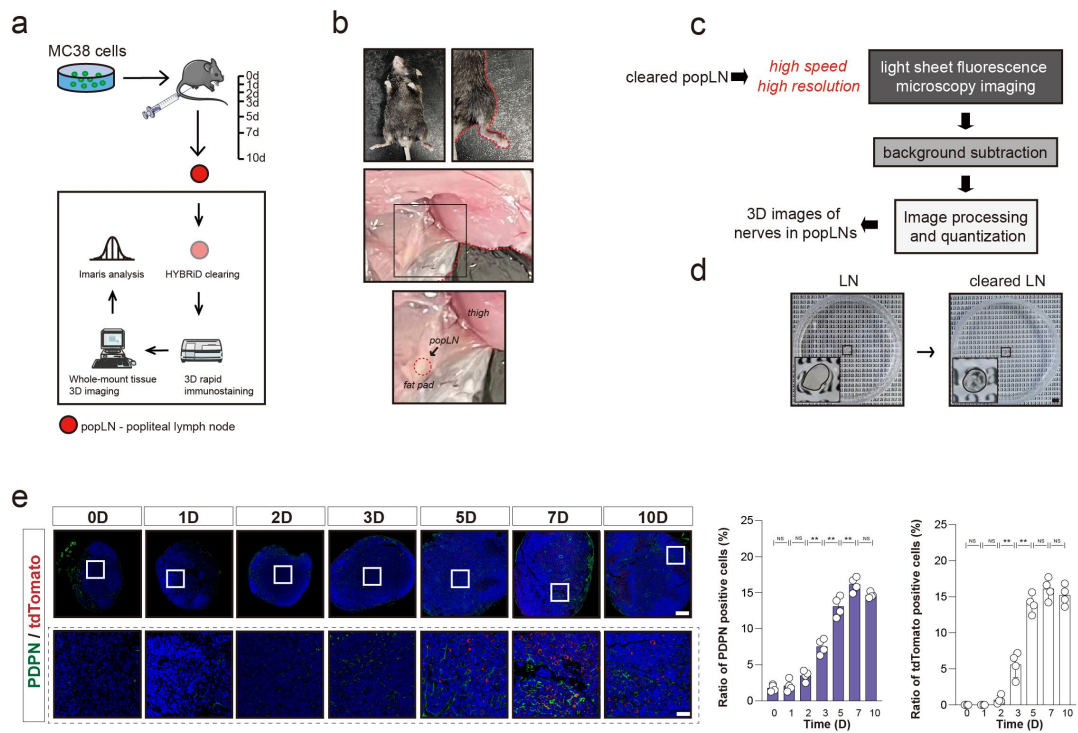

**Extended Data Figure 1. Experimental workflow and time-course analysis of tumor-induced**

**popLN remodeling.**

a. The popLN metastasis model was established using MC38 colorectal cancer cells that were

injected into the footpads of C57BL/6 mice.

b. The location of popLNs in C57BL/6 mice.

c. Schematic of the whole-organ clearing, imaging, and image-analysis workflow.

d. Photos of popLN tissues before and after HYBRiD clearing. Scale bars: 3 mm.

e. Immunofluorescent staining of tdTomato/PDPN/DAPI in popLNs at days 0, 1, 2, 3, 5, 7, and 10.

Scale bars: upper - 300 μm; lower - 100 μm.

\*\*: P<0.01; NS: no significant difference.

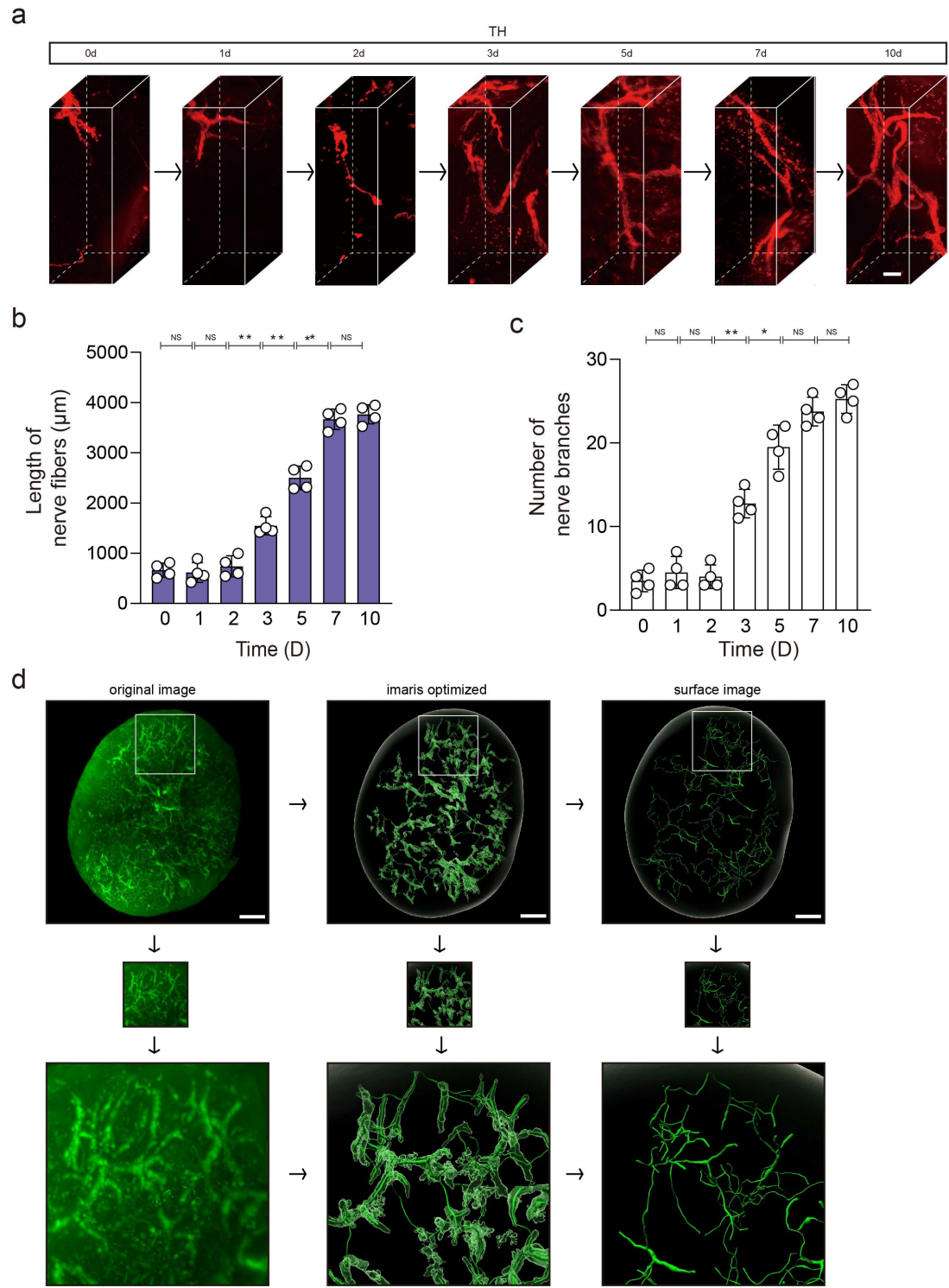

**Extended Data Figure 2. 3D images of the sympathetic nerves in popLNs at different time points.**

a-c. 3D images of TH<sup>+</sup> sympathetic nerve fibers in popLNs at days 0, 1, 2, 3, 5, 7, and 10 (a). A representative 3D volume (420 μm × 1,000 μm × 420 μm) is shown. Quantification of the length of nerve fibers (b) and the number of nerve branches (c) are shown below. Scale bars: 150 μm.

d. 3D images of the original and surface images of popLN obtained at day 7. The images were

processed using Imaris. Scale bars: 300 μm.

\*: $P < 0.05$; NS: no significant difference.

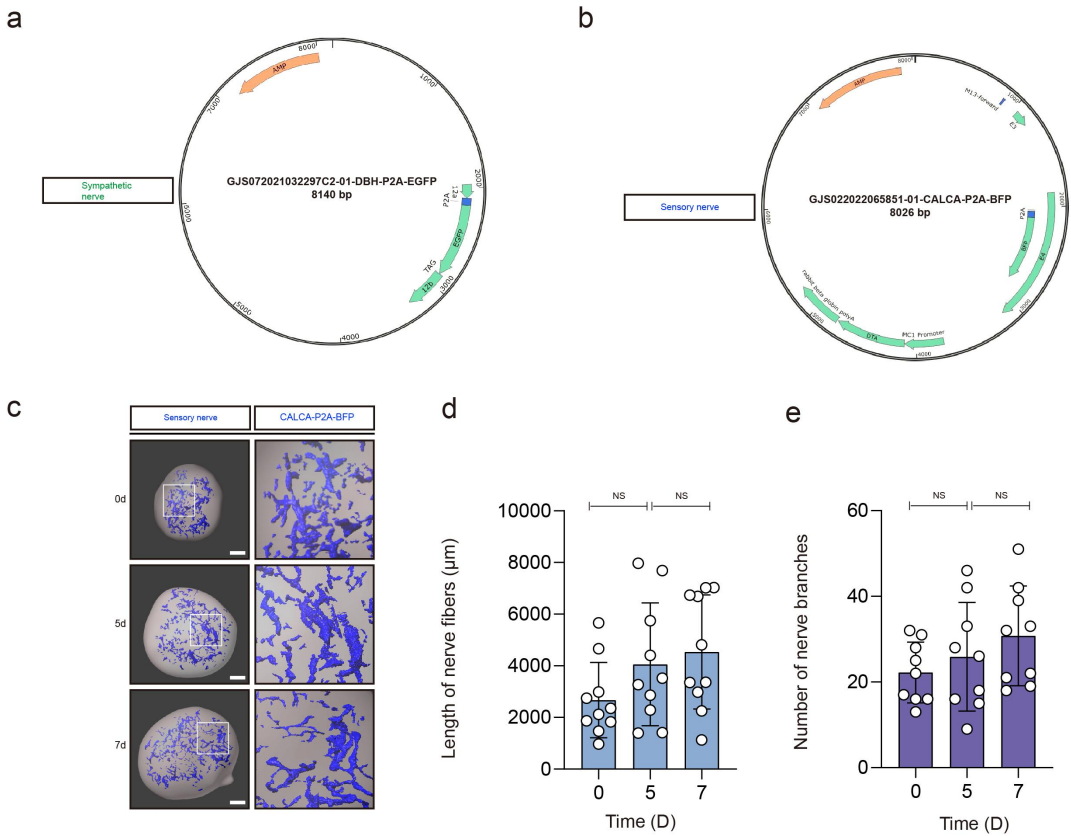

**Extended Data Figure 3. Generation of reporter mice and analysis of sensory innervation in**
**popLNs.**

a, b. The structure of the targeting vectors (DBH-P2A-EGFP, a; CALCA-P2A-BFP, b).

c-e. 3D images of the sensory nerves in popLNs at days 0, 5 and 7. CALCA-BFP<sup>+</sup> sensory nerve
fibers are shown in blue (c). Quantification of the length of nerve fibers (d) and the number of
nerve branches (e) are shown on the right. Scale bars: 300 μm.

NS: no significant difference.

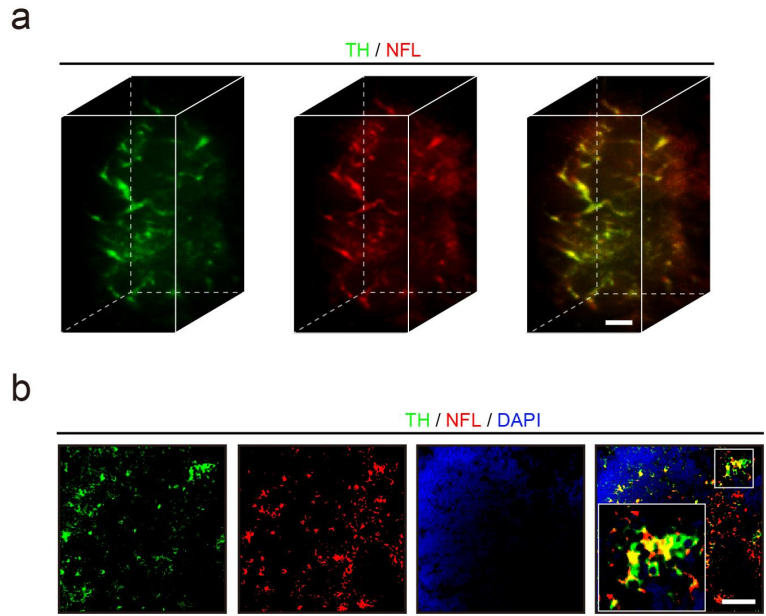

**Extended Data Figure 4. 3D images and immunofluorescent staining of NFL and TH in popLNs.**

a. 3D images of NFL and TH in popLN at day 7. Scale bars: 100 μm.

b. Immunofluorescent staining of NFL and TH in popLN at day 7. Scale bars: 100 μm.

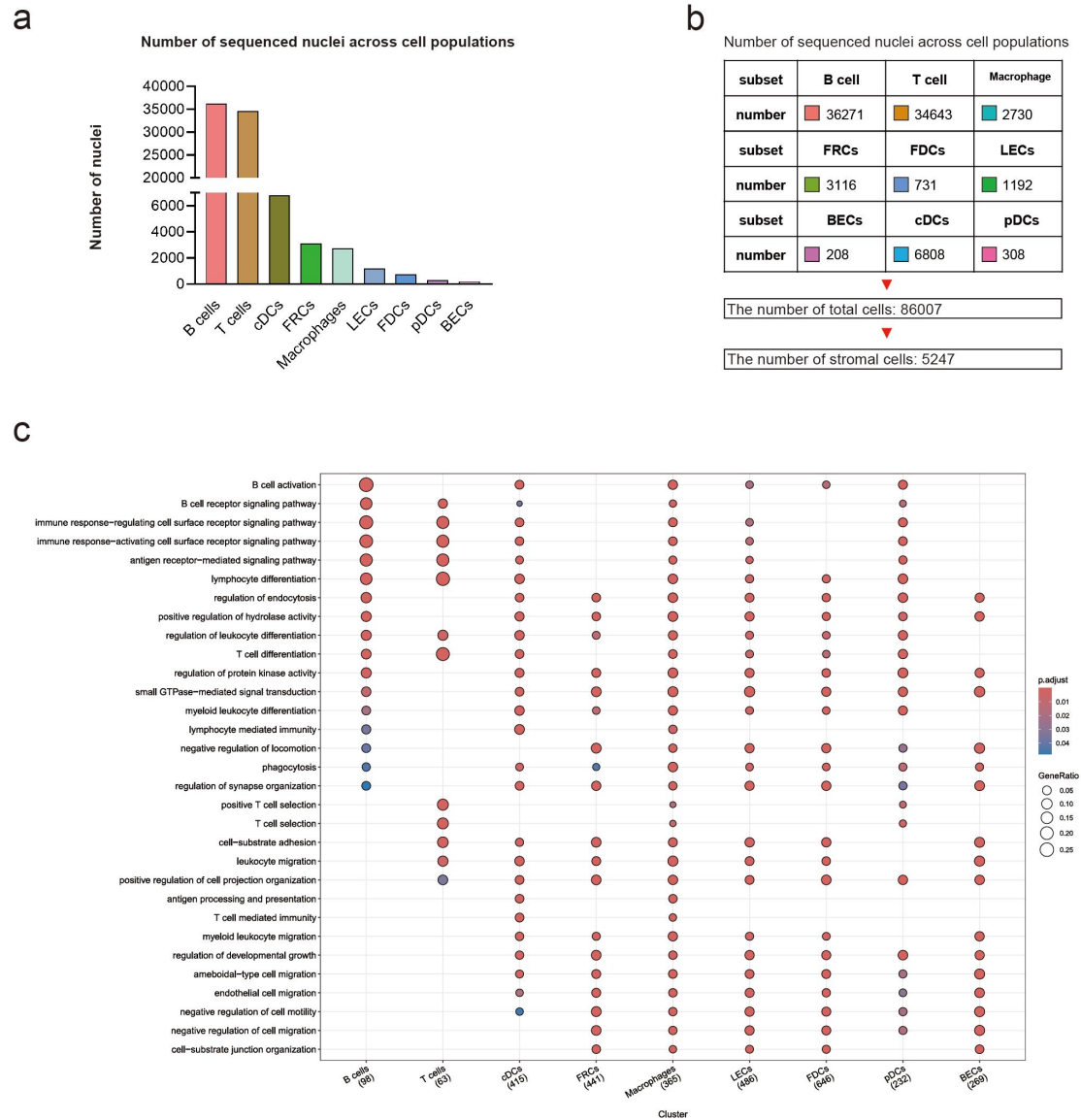

**Extended Data Figure 5. snRNA-seq analyses of different cell subsets in popLNs.**

a, b. Distribution of sequenced nuclei across the indicated cell populations.

c. GO-BP analysis of different cell subsets in the popLNs from days 0 to 10. The dataset has been deposited in the Gene Expression Omnibus under accession number GSE280461.

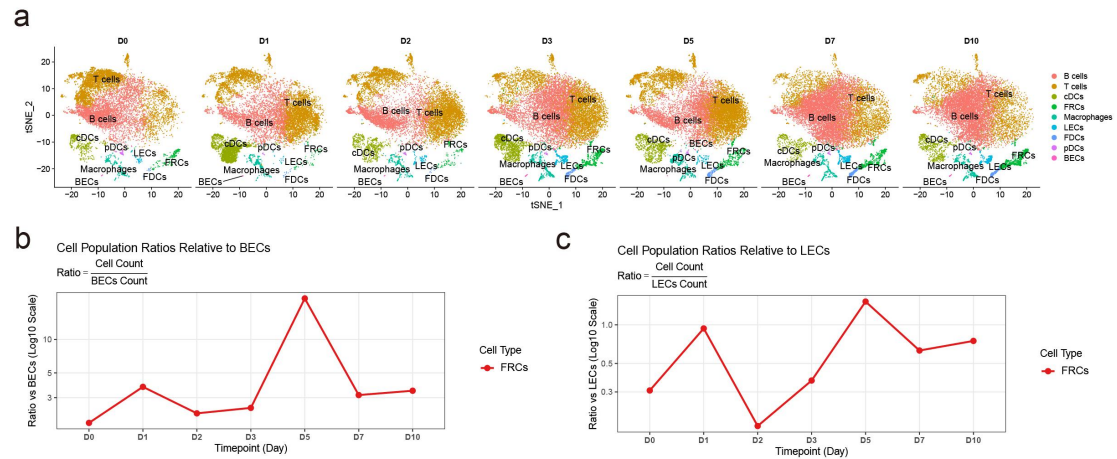

**Extended Data Figure 6. snRNA-seq analyses of different cell subsets at different time points.**

a. t-SNE visualization of different cell subsets in popLNs at days 0, 1, 2, 3, 5, 7, and 10.

b, c. Ratios of FRC nuclei counts to BEC nuclei counts (b) and LEC nuclei counts (c) at the indicated time points.

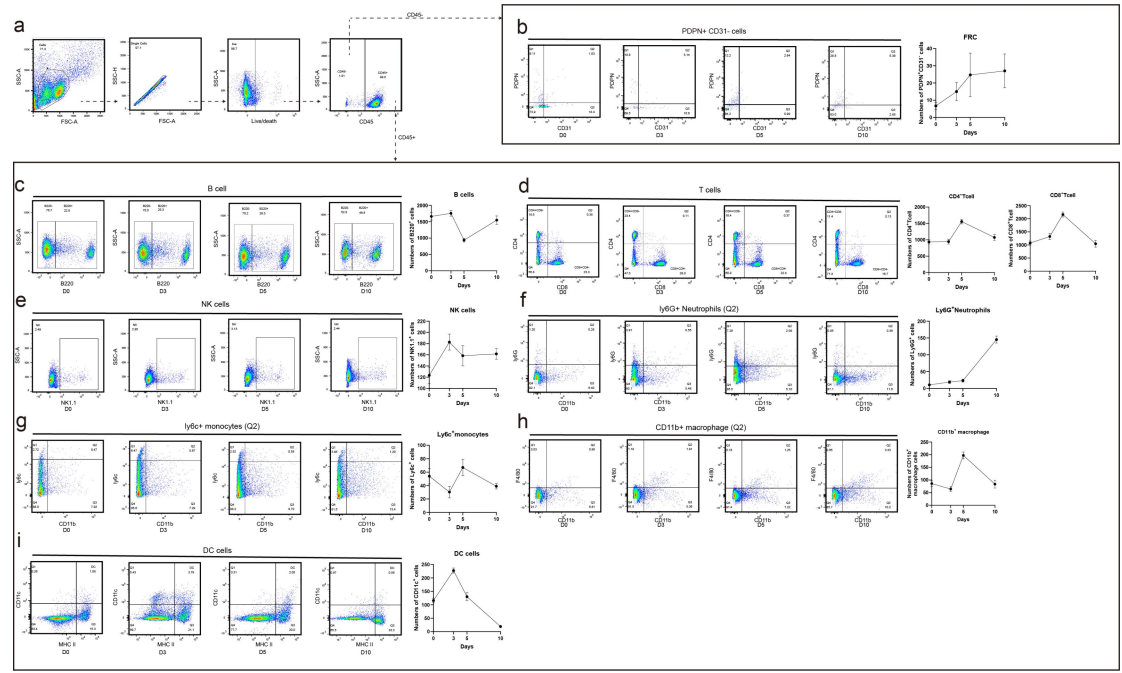

**Extended Data Figure 7. Changes in stromal and immune cell populations in popLNs during tumor-induced LN remodeling.**

a. Gating strategy for flow cytometric analysis. Cells were sequentially gated as FSC/SSC, singlets, live cells, and CD45-positive or CD45-negative populations for downstream analysis.

b. Representative flow cytometry plots and normalized cell numbers of PDPN<sup>+</sup>CD31<sup>-</sup> stromal cells within the CD45<sup>-</sup> compartment at the indicated time points.

c-i. Representative flow cytometry plots and normalized cell numbers of B cells (c), CD4<sup>+</sup> T cells and CD8<sup>+</sup> T cells (d), NK cells (e), Ly6G<sup>+</sup> neutrophils (f), Ly6C<sup>+</sup> monocytes (g), CD11b<sup>+</sup>F4/80<sup>+</sup> macrophages (h), and dendritic cells (DCs) (i). Each group was repeated at least 3 times.

Cell numbers were normalized to $1 \times 10^4$ viable cells per popLN and should not be interpreted as absolute cell numbers per lymph node.

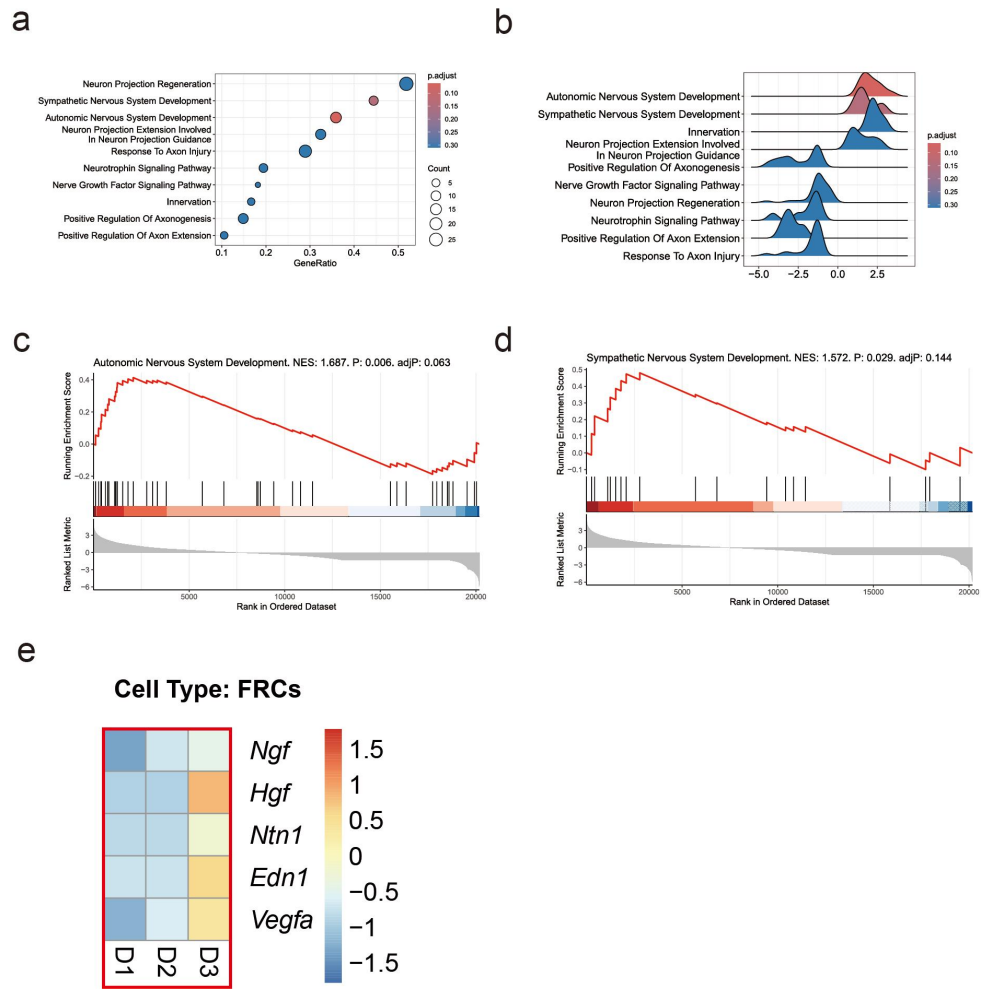

**Extended Data Figure 8. GO and GSEA analyses of neural-related transcriptional programs in FRCs.**

a, b. GO analysis of neural-related biological processes enriched in FRCs.

c, d. GSEA of autonomic nervous system development and sympathetic nervous system development signatures in FRCs.

e. Heatmap of *Ngf*, *Hgf*, *Ntn1*, *Edn1* and *Vegfa* at days 1, 2 and 3.

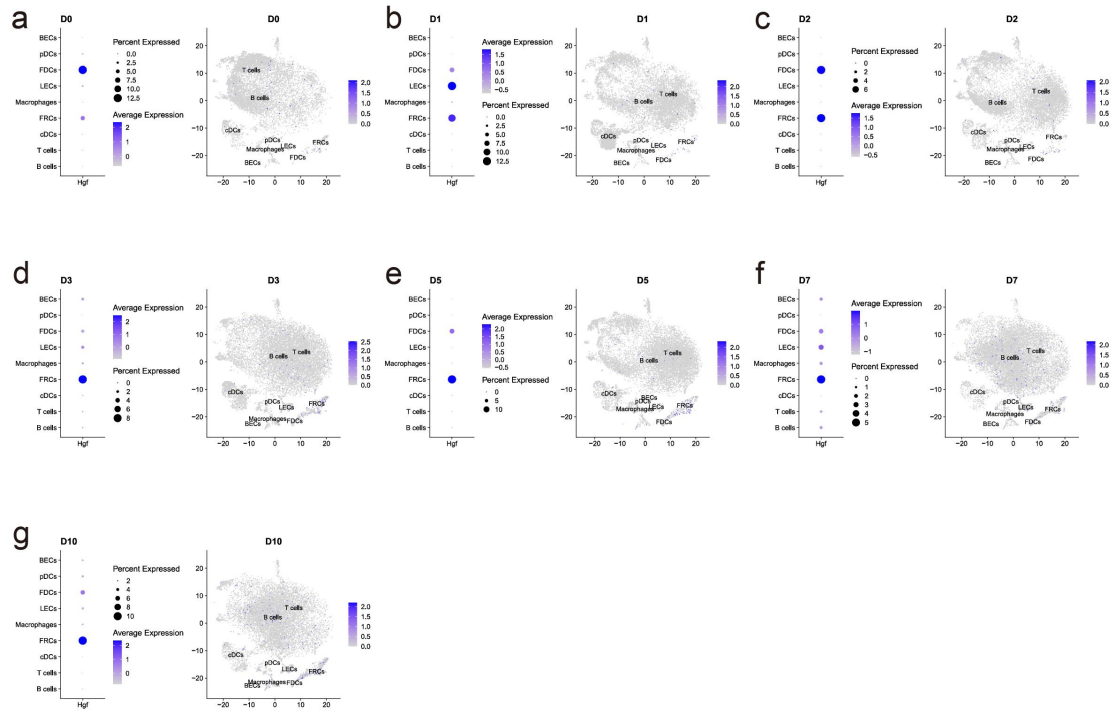

70

71 **Extended Data Figure 9. snRNA-seq analyses of *Hgf* expression at different time points.**

72 a-g. The expression of *Hgf* in different cell subsets is displayed by t-SNE visualization at days 0

73 (a), 1 (b), 2 (c), 3 (d), 5 (e), 7 (f), and 10 (g).

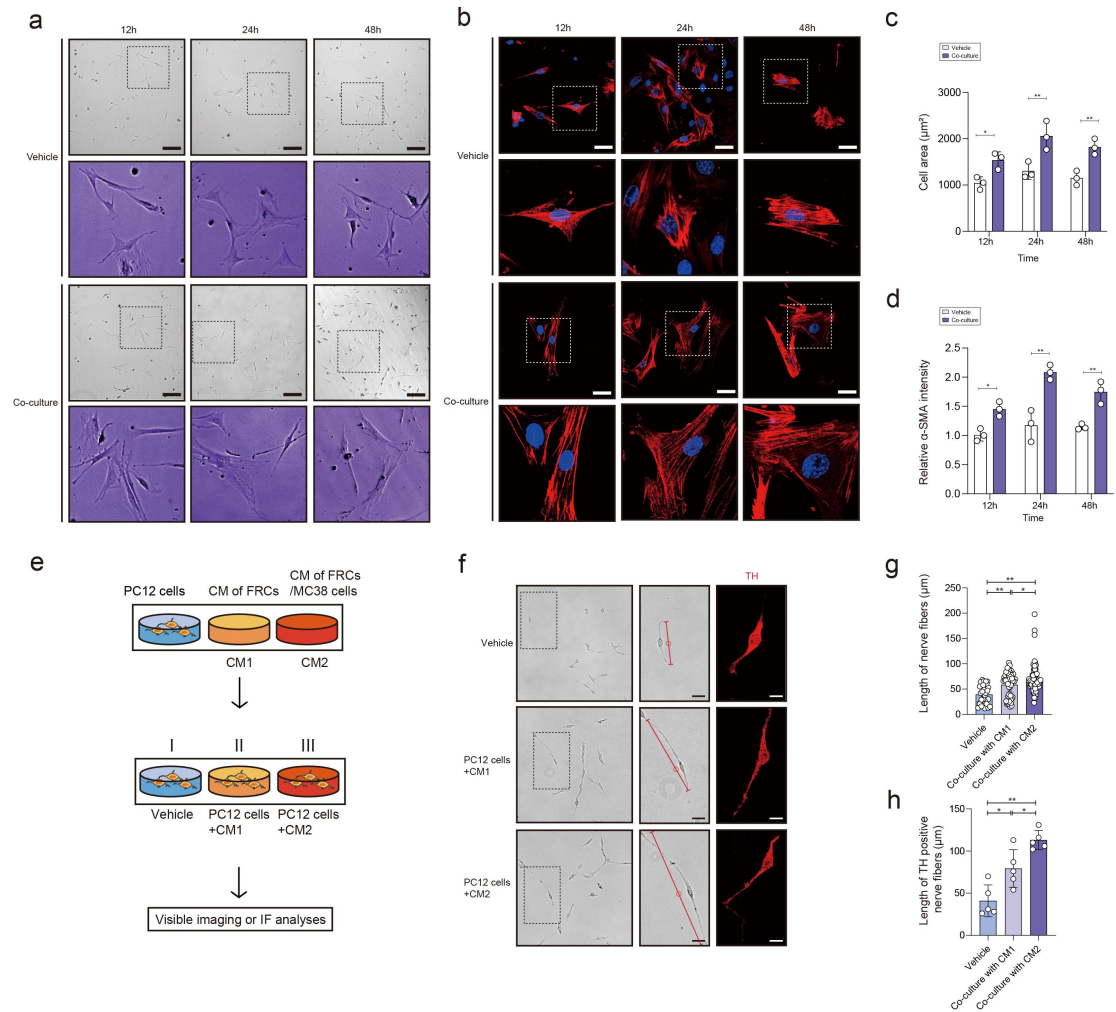

**Extended Data Figure 10. Responses of primary FRCs and PC12 cells to MC38-conditioned medium and FRC-conditioned medium.**

a. Representative bright-field images of primary FRCs treated with MC38-conditioned medium or the control at 12, 24 and 48 h. Scale bars: 40 μm.

b. Immunofluorescent staining of α-SMA in primary FRCs treated with MC38-conditioned medium or the control at 12, 24 and 48 h. Scale bars: 10 μm.

c, d. Quantification of cell area of primary FRCs (c) and relative α-SMA intensity in primary FRCs (d) under the indicated conditions at 12, 24 and 48 h.

e. Schematic of PC12 cells that were treated with the indicated FRC-conditioned media (CM1: conditioned medium from FRCs alone; CM2: conditioned medium from FRCs exposed to MC38-conditioned medium).

f-h. Representative bright-field and immunofluorescent images of PC12 cells under the indicated conditions at 48 h (f). Quantification of the length of nerve fibers in the bright-field (g) and immunofluorescent images (h) are shown on the right. Scale bars, 20 μm (left) and 12 μm (right).

For bright-field quantification, all PC12 cells within each analyzed field were included; for immunofluorescence-based quantification, at least five PC12 cells were analyzed per group. Each experiment was repeated at least five times.

\*: P<0.05; \*\*: P<0.01.

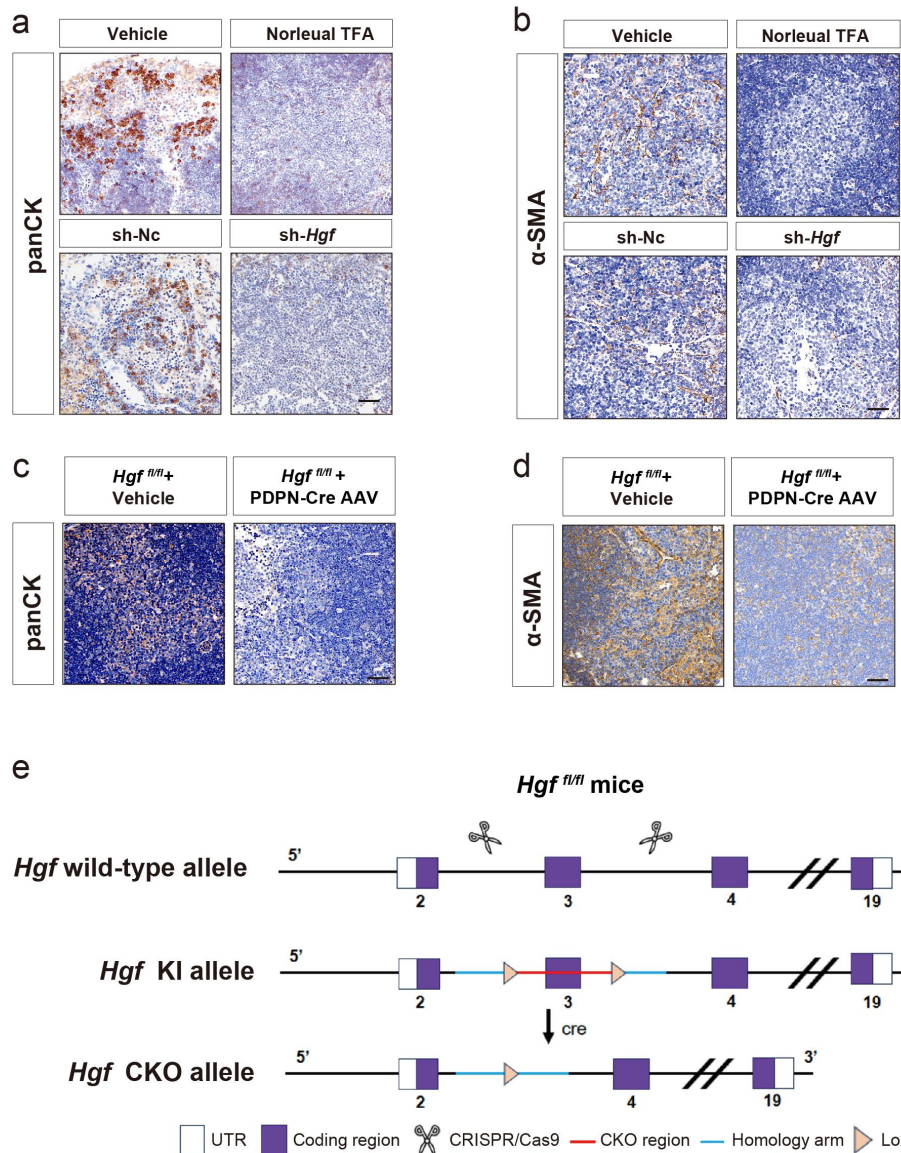

**Extended Data Figure 11. Immunohistochemical staining of panCK and α-SMA in popLNs.**

a, b. Immunohistochemical staining of panCK (a) and α-SMA (b) in popLNs treated with HGF inhibitor or vehicle, AAV-sh-*Hgf* or sh-NC and inoculated with MC38 cells after 7 days. Scale bars: 100 μm.

c, d. Immunohistochemical staining of panCK (c) and α-SMA (d) in popLNs treated with PDPN-Cre AAV or vehicle and inoculated with MC38 cells after 7 days. Scale bars: 100 μm.

e. Schematic of the construction of *Hgf*<sup>fl/fl</sup> mouse.

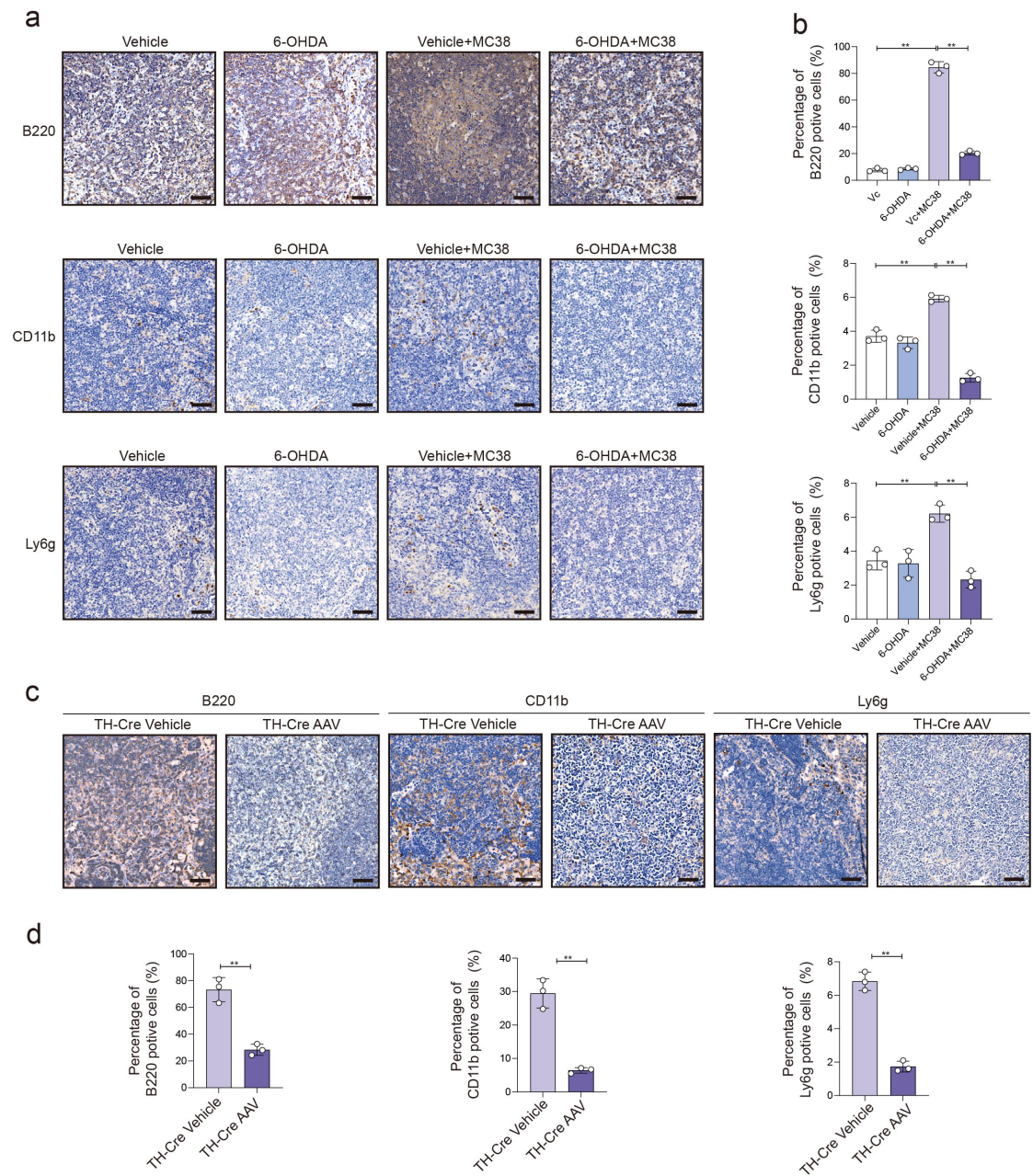

**Extended Data Figure 12. Effects of local sympathetic denervation on B220<sup>+</sup> B cells, CD11b<sup>+</sup> myeloid cells, and Ly6G<sup>+</sup> neutrophils in popLNs.**

a. Immunohistochemical staining of B220, CD11b and Ly6G in popLNs treated with 6-OHDA or vehicle and inoculated with MC38-tdTomato cells after 7 days. Scale bars: 100 μm.

b. Quantification of the percentages of B220, CD11b and Ly6G positive cells in popLNs under the indicated conditions.

c. Immunohistochemical staining of B220, CD11b and Ly6G in popLNs from TH-Cre mice treated with rAAV or vehicle and inoculated with MC38-tdTomato cells after 7 days. Scale bars: 100 μm.

d. Quantification of the percentages of B220, CD11b and Ly6G positive cells in popLNs under the indicated conditions.

\*\*: P<0.01.

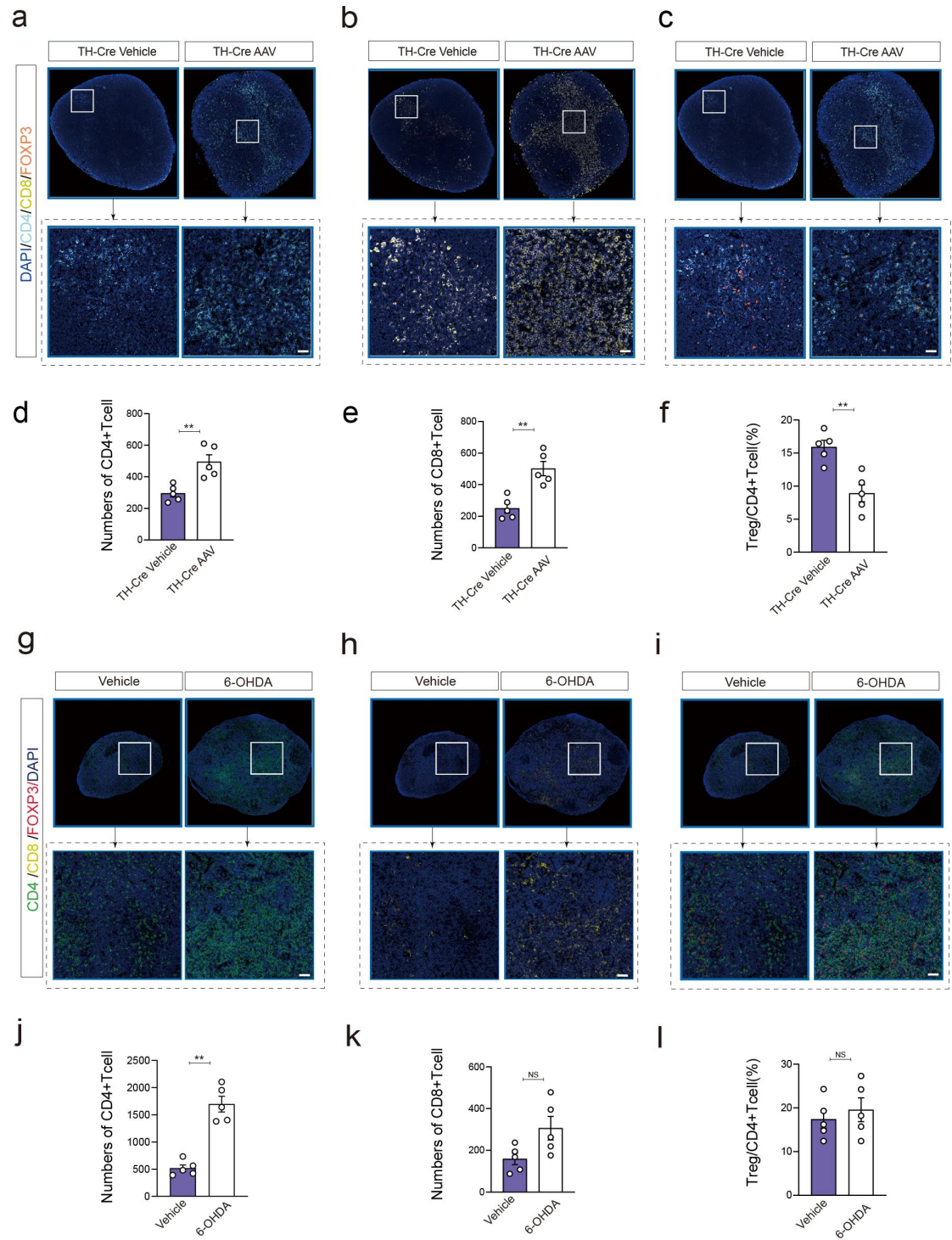

**Extended Data Figure 13. Immunofluorescent staining of CD4<sup>+</sup> T cells, CD8<sup>+</sup> T cells and Treg/CD4<sup>+</sup> T-cell ratio in popLNs treated with 6-OHDA or rAAV.**

a-f. Immunofluorescent staining of CD4 (a), CD8 (b) and FOXP3/CD4 (c) in popLNs treated with rAAV or vehicle and inoculated with MC38-tdTomato cells after 7 days. Quantification of the number of CD4<sup>+</sup> T cells (d), CD8<sup>+</sup> T cells (e) and Treg/CD4<sup>+</sup> T-cell ratio (f) in popLNs are shown below. Scale bars: 100 μm.

g-l. Immunofluorescent staining of CD4 (g), CD8 (h) and FOXP3/CD4 (i) in popLNs treated with 6-OHDA or vehicle and inoculated with MC38-tdTomato cells after 7 days. Quantification of the number of CD4<sup>+</sup> T cells (j), CD8<sup>+</sup> T cells (k) and Treg/CD4<sup>+</sup> T-cell ratio (l) in popLNs are shown below. Scale bars: 100 μm.

\*\*<sub>2</sub>: P<0.01.

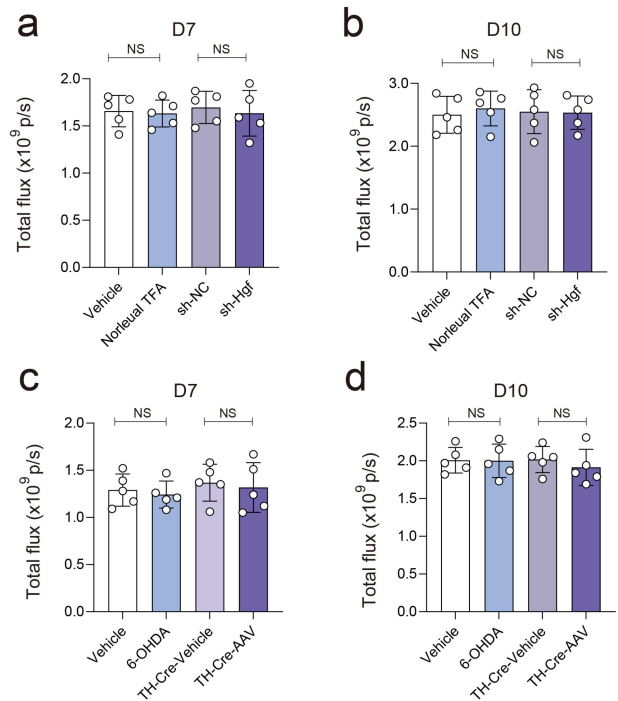

**Extended Data Figure 14. Effects of local HGF targeting and sympathetic denervation on primary tumor burden.**

a, b, Primary footpad tumor burden, quantified by tdTomato radiant efficiency, at days 7 (a) and 10 (b) in C57BL/6 mice treated with Norleual TFA, AAV-sh-Hgf, or the corresponding controls.

c, d, Primary footpad tumor burden, quantified by tdTomato radiant efficiency, at days 7 (c) and 10 (d) following local 6-OHDA-mediated sympathetic denervation or TH-Cre-dependent taCasp3-mediated genetic sympathetic ablation, together with the corresponding controls.

NS: no significant difference.

**Extended Data Video 1. 3D videos of DBH-EGFP<sup>+</sup> nerve fibers in popLNs at days 0, 1, 2, 3, 5, 7, and 10.**

The time periods during which popLNs at each time point appear in the video: 0-3 seconds – day 0; 4-7 seconds – day 1; 8-11 seconds – day 2; 12-16 seconds – day 3; 17-22 seconds – day 5; 23-29 seconds – day 7; 30-36 seconds – day 10.

| Target | Fluorophore / enzyme | Host | Clone | Company | Catalogue no. | Application | Working dilution / concentration |
| --- | --- | --- | --- | --- | --- | --- | --- |
| α-SMA (ACTA2) | Unconjugated | Rabbit | EPR5368 | Abcam | ab124964 | Cell IF; DAB-IHC-P | 1:400 (cell IF); 1:2,000 (IHC-P) |
| CD4 | FITC | Rat | GK1.5 | BioLegend | 100405 | Flow cytometry | 1:400 |
|  | Unconjugated | Rabbit | EPR19514 | Abcam | ab183685 | Tissue IF (CD4–FOXP3) | 1:1,000 |
| CD8a | Brilliant Violet 510 | Rat | 53-6.7 | BioLegend | 100752 | Flow cytometry | 1:400 |
|  | Unconjugated | Rat | 4SM15 | Invitrogen / eBioscience | 14-0808-82 | Tissue IF | 1:1,000 |
| CD11b | BUV805 | Rat | M1/70 | Invitrogen / eBioscience | 368-0112-82 | Flow cytometry | 1:400 |
|  | Unconjugated | Rabbit | EPR1344 | Abcam | ab133357 | DAB-IHC-P | 1:4,000 |
| CD11c | PE | Armenian hamster | N418 | BioLegend | 117308 | Flow cytometry | 1:400 |
| CD31 | APC | Rat | 390 | BioLegend | 102510 | Flow cytometry | 1:400 |
| CD45 | PE | Rat | 30-F11 | BioLegend | 103106 | Flow cytometry | 1:400 |
|  | PerCP | Rat | 30-F11 | BioLegend | 103130 | Flow cytometry | 1:400 |
| CD45R / B220 | Brilliant Violet 421 | Rat | RA3-6B2 | BioLegend | 103251 | Flow cytometry | 1:400 |
|  | Unconjugated | Rat | RA3-6B2 | Invitrogen / eBioscience | 14-0452-82 | DAB-IHC-P | 1:1,000 |
| F4/80 | BUV661 | Rat | T45-2342 | BD Biosciences | 750643 | Flow cytometry | 1:400 |
| FOXP3 | APC | Rat | FJK-16s | Invitrogen / eBioscience | 17-5773-82 | Flow cytometry | 1:400 |
|  | Unconjugated | Rat | FJK-16s | Invitrogen / eBioscience | 14-5773-82 | Tissue IF (CD4–FOXP3) | 1:1,000 |
| HGF (HGFα) | Alexa Fluor 647 | Mouse | H-10 | Santa Cruz Biotechnology | sc-374422 AF647 | Tissue IF (PDPN–HGF) | 1:200 |
| I-A/I-E (MHC II) | PE/Cyanine7 | Rat | M5/114.15.2 | BioLegend | 107630 | Flow cytometry | 1:1,000 |
| Ly-6C | FITC | Rat | HK1.4 | BioLegend | 128006 | Flow cytometry | 1:1,000 |
| Ly-6G | PE | Rat | 1A8 | BioLegend | 127607 | Flow cytometry | 1:1,000 |
|  | Unconjugated | Rabbit | EPR22909-135 | Abcam | ab238132 | DAB-IHC-P | 1:2,000 |
| MET (c-Met) | Unconjugated | Rabbit | Polyclonal | Invitrogen | PA5-85951 | Tissue IF (MET–PGP9.5 ; MET–TH) | 1:200 |
|  | Unconjugated | Rabbit | EPR22436-24 | Abcam | ab216330 | Cell IF (MET–TH) | 1:50 |
| NFL (NEFL) | Unconjugated | Rabbit | T.400.5 | Invitrogen | MA5-14981 | Whole-mount IF; tissue IF (TH–NFL) | 1:100 (whole-mount IF); 1:150 (tissue IF) |
| NK-1.1 | PE/Cyanine7 | Mouse | PK136 | BioLegend | 108714 | Flow cytometry | 1:400 |
| pan Cytokeratin | Unconjugated | Mouse | C-11 | Abcam | ab7753 | Mouse-on-mouse DAB-IHC-P | 1:1,000 |
| M.O.M. ImmPRESS anti-Mouse IgG HRP polymer | HRP | Horse | Polyclonal polymer | Vector Laboratories | MP-2400 | Mouse-on-mouse DAB-IHC-P (pan Cytokeratin) | Ready-to-use |
| Podoplanin (PDPN) | Alexa Fluor 488 | Syrian hamster | eBio8.1.1 (8.1.1) | Invitrogen / eBioscience | 53-5381-82 | Whole-mount IF; tissue IF (PDPN–HGF) | 1:1,000 |
|  | PE | Syrian hamster | 8.1.1 | BioLegend | 127408 | Flow cytometry | 1:400 |
| PGP9.5 (UCHL1) | Unconjugated | Rabbit | Polyclonal | Invitrogen | PA5-29012 | Whole-mount IF | 1:100 |
|  | Unconjugated | Chicken | Polyclonal | Novus Biologicals / Bio-Techne | NB110-58872 | Tissue IF (PGP9.5–MET) | 1:2,000 |
| TH | Unconjugated | Rabbit | Polyclonal | Abcam | ab112 | Whole-mount IF | 1:100 |
|  | Unconjugated | Chicken | Polyclonal | Abcam | ab76442 | Tissue IF (TH–NFL; TH–MET) | 1:1,500 |
|  | Unconjugated | Chicken | Polyclonal | Invitrogen | PA5-143583 | Cell IF (TH–βIII-tubulin; TH–MET) | 1:2,000 |
| Donkey anti-Mouse IgG (H+L) | Alexa Fluor 488 | Donkey | Polyclonal | Jackson ImmunoResearch | 715-545-150 | Cell IF; tissue IF; whole-mount IF | 1:500 |
|  | Alexa Fluor 594 | Donkey | Polyclonal | Jackson ImmunoResearch | 715-585-150 | Cell IF; tissue IF; whole-mount IF | 1:500 |
|  | Alexa Fluor 647 | Donkey | Polyclonal | Jackson ImmunoResearch | 715-605-150 | Cell IF; tissue IF; whole-mount IF | 1:500 |
| Donkey anti-Rabbit IgG (H+L) | Alexa Fluor 488 | Donkey | Polyclonal | Jackson ImmunoResearch | 711-545-152 | Cell IF; tissue IF; whole-mount IF | 1:500 |
|  | Alexa Fluor 594 | Donkey | Polyclonal | Jackson ImmunoResearch | 711-585-152 | Cell IF; tissue IF; whole-mount IF | 1:500 |
|  | Alexa Fluor 647 | Donkey | Polyclonal | Jackson ImmunoResearch | 711-605-152 | Cell IF; tissue IF; whole-mount IF | 1:500 |
| Donkey anti-Chicken IgY (H+L) | Alexa Fluor 488 | Donkey | Polyclonal | Jackson ImmunoResearch | 703-545-155 | Tissue IF; cell IF | 1:500 |
|  | Alexa Fluor 594 | Donkey | Polyclonal | Jackson ImmunoResearch | 703-585-155 | Tissue IF; cell IF | 1:500 |
| Donkey anti-Rat IgG (H+L) | Alexa Fluor 488 | Donkey | Polyclonal | Jackson ImmunoResearch | 712-545-153 | Tissue IF | 1:500 |
|  | Alexa Fluor 594 | Donkey | Polyclonal | Jackson ImmunoResearch | 712-585-153 | Tissue IF | 1:500 |
|  | Alexa Fluor 647 | Donkey | Polyclonal | Jackson<br>ImmunoRese<br>arch | 712-605-15<br>3 | Tissue IF | 1:500 |
| Donkey<br>anti-Rab<br>bit IgG<br>(H+L) | HRP | Donkey | Polyclonal | Jackson<br>ImmunoRese<br>arch | 711-035-15<br>2 | DAB-IHC-P<br>(α-SMA;<br>CD11b; Ly-6G) | 1:2,500 |
| Donkey<br>anti-Rat<br>IgG<br>(H+L) | HRP | Donkey | Polyclonal | Jackson<br>ImmunoRese<br>arch | 712-035-15<br>3 | DAB-IHC-P<br>(CD45R/B220) | 1:2,500 |

142 **Extended Data Table 1. Antibody list.**

143
